# PP2A and GSK3 act as modifiers of FUS-ALS by modulating mitochondrial transport

**DOI:** 10.1101/2023.08.28.555106

**Authors:** Paraskevi Tziortzouda, Jolien Steyaert, Wendy Scheveneels, Adria Sicart, Katarina Stoklund Dittlau, Adriana Margarida Barbosa Correia, Arun Pal, Andreas Hermann, Philip Van Damme, Thomas Moens, Ludo Van Den Bosch

## Abstract

ALS is a fatal neurodegenerative disease which currently lacks effective treatments. Mutations in the RNA-binding protein FUS are a common cause of familial ALS, accounting for around 4% of fALS cases. Studying the mechanisms by which mutant FUS is toxic to neurons may provide insight into the pathogenesis of both familial and sporadic forms of ALS. Here we identify Protein Phosphatase 2A (PP2A) and Glycogen Synthase Kinase 3 (GSK3) as novel modifiers of FUS-ALS *in vivo*, looking from fly to human. PP2A-C and GSK3β inhibition rescued FUS-induced toxicity in *Drosophila* and disease-relevant phenotypes in human iPSC-derived spinal motor neurons (sMNs). In both *Drosophila* and human iPSC-sMNs, we observed reduced GSK3β inhibitory phosphorylation, suggesting that FUS dysfunction results in GSK3β hyperactivity. We found that PP2A acts upstream of GSK3, affecting its inhibitory phosphorylation, and in synergy they modulate mitochondrial transport through the motor protein kinesin. Our data provide *in vivo* evidence that PP2A and GSK3 are disease modifiers, and reveal an unexplored mechanistic link between PP2A, GSK3 and kinesin in FUS-associated ALS.

## Introduction

Amyotrophic lateral sclerosis (ALS) is the most common motor neuron disease in adults, characterized by selective loss of upper and lower motor neurons in the brain and spinal cord^1,2^. Symptoms include muscle wasting and paralysis, and death usually occurs within an average of three years post-diagnosis due to failure of the respiratory muscles^2^. In 90% of cases ALS is sporadic (sALS) with no known family history^1^. However, 10% of cases are familial (fALS), where patients inherit the disease mostly in an autosomal dominant manner^1^. Mutations in the *Fused in Sarcoma (FUS)* gene, encoding for FUS protein, account for ∼4% of fALS and ∼1% of sALS cases^1^. FUS is a DNA/RNA-binding protein, and in mutation carriers, it mislocalizes to the cytoplasm where it aggregates^1^.

Despite advances in the understanding of the genetics and pathogenesis of ALS, effective therapeutic options are currently lacking, with licensed drugs extending the lifespan of patients by only a few months^2^. Here we aimed to identify new targets involved in the pathogenesis of FUS-associated ALS using *Drosophila* and induced pluripotent stem cell-derived spinal motor neuron (iPSC-sMN) models.

As a human model of FUS-associated fALS, we previously generated and extensively characterized iPSC-sMNs from *FUS* mutation carriers as well as their isogenic controls^3–5^. In these sMNs we observed FUS cytoplasmic mislocalization, similar to that reported in cellular models and patient post mortem material^6^. In addition, we observed a mitochondrial neuritic transport defect^3^, which we have since confirmed is a common phenotype in iPSC-derived sMNs from *TARDBP* and *C9orf72* familial-ALS mutation carriers^7,8^. This suggests that mitochondrial transport defects could be a common underlying disease mechanism in ALS. Recently, we have also demonstrated that the *FUS* mutant sMNs fail to correctly form neuromuscular junctions (NMJs) when cultured with human primary mesoangioblast-derived myotubes in microfluidic devices^4^.

We previously showed that overexpression of wild-type or ALS-mutant FUS in *Drosophila* MNs is toxic, leading flies to die in their pupal cases. Therefore, this *Drosophila* model provides a useful screening tool to identify candidate modifying genes^9,10^. Using eclosion as a readout, we performed a genome-wide genetic screen to identify modifiers of FUS-ALS *in vivo*, and found that *microtubule star* (*mts*) (the *Drosophila* ortholog of *PP2A-C*) and *shaggy* (*sgg*) (the *Drosophila* ortholog of *GSK3β*), are novel modifiers of FUS-associated ALS. Specifically, genetic and pharmacological inhibition of mts and sgg rescued eclosion and lifespan of FUS flies. Importantly, PP2A and GSK3β pharmacological inhibition in human cells rescued hallmark ALS-associated phenotypes, including FUS cytoplasmic mislocalization, NMJ formation and mitochondrial transport defects. Interestingly, GSK3 inhibitory phosphorylation appeared reduced in our FUS-ALS models, suggesting that FUS dysfunction results in GSK3 hyperactivity. We also found that the phosphatase PP2A acts upstream of GSK3, altering its inhibitory phosphorylation, in both flies and human cells.

Mitochondrial transport is a vital process mainly mediated by the motor protein kinesin^11,12^, and GSK3β has been shown to phosphorylate kinesin-1 in *Drosophila* and mammalian cells^13,14^. Hence, GSK3β hyperactivity could cause kinesin hyperphosphorylation and dysfunction, explaining the axonal transport deficits observed in patient sMNs. Excitingly, we observed that PP2A and GSK3 inhibition can rescue mitochondrial transport deficits in FUS-ALS patient iPSC-sMNs, and kinesin-1 appears as an intermediate regulator of this cascade. Altogether, by looking from fly to human, our data provide further insight into the mechanisms of FUS toxicity, and have identified PP2A-C and GSK3β as novel disease modifiers and potential therapeutic targets for ALS.

## Results

### A genome-wide screen identifies *mts* and *sgg* as candidate modifiers of FUS-ALS *in vivo*

We performed a genome-wide screen to identify candidate modifiers (suppressors) of FUS toxicity in *Drosophila* (Fig. 1A). We previously showed that motor neuron specific expression of wild-type or mutant human FUS leads to a severe eclosion phenotype^9^. We therefore generated a recombinant stock of UAS-FUS (R521G) and the D42-Gal4 motor neuron driver line. This stock was maintained over a TM6B-Gal80 balancer, which suppresses expression and is marked with a visible marker. We crossed this screening stock to flies from Bloomington *Drosophila* Stock Center (BDSC) deficiency kit, which consists of a selected set of molecularly defined genomic deletions. We focused on deletions on chromosomes X/Y, 2 and 3, crossing 473 deficiency lines to the screening stock and assessing whether eclosion was rescued. When crossed to a control background, the D42Gal4>FUS(R521G) flies are not able to eclose, therefore any eclosion was considered as a rescue. Using this approach, we identified 58 candidate deficiencies that attenuated the mutant FUS pupal lethality.

**Fig. 1.**
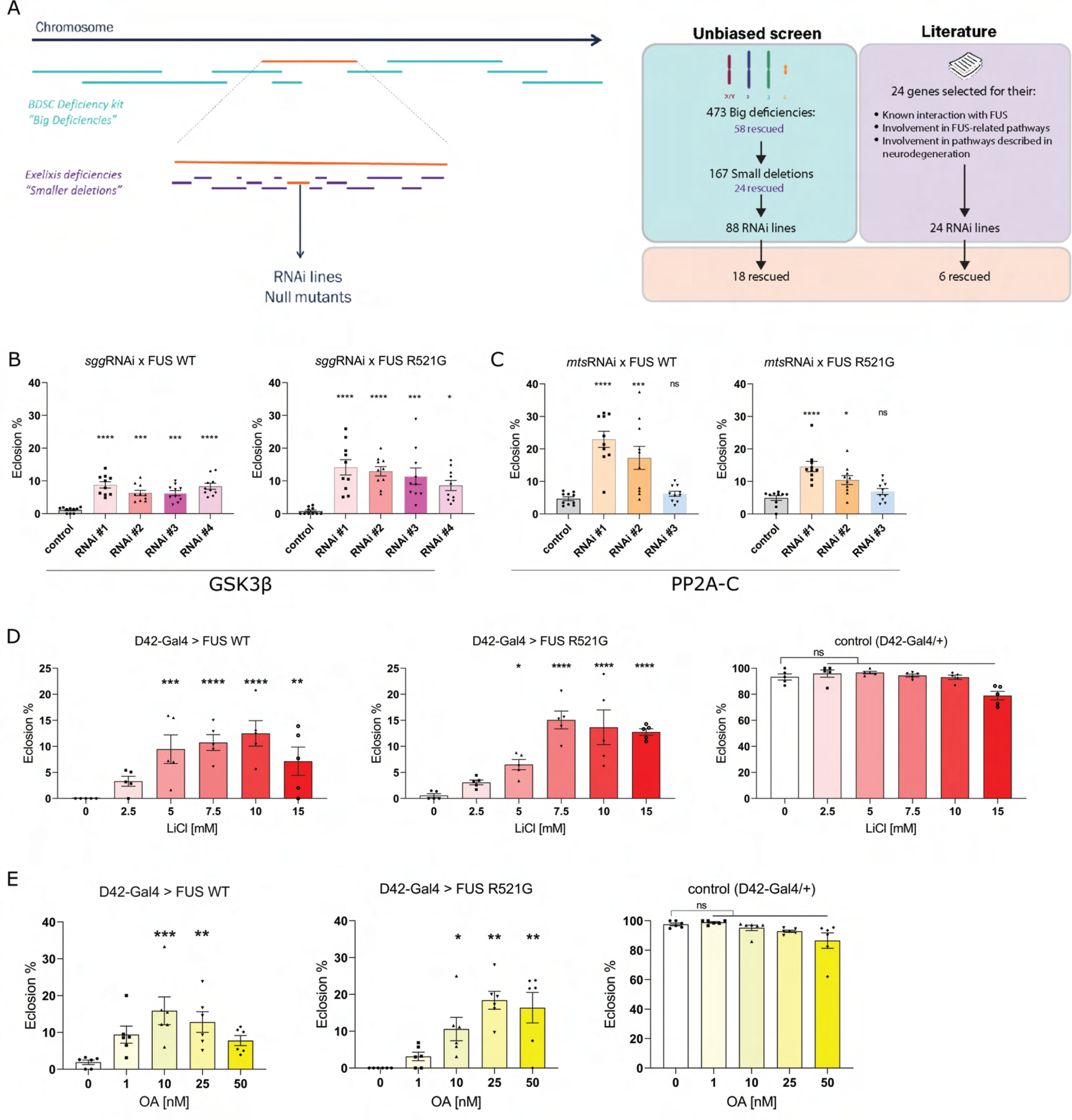
*mts* and *sgg* are modifiers of FUS toxicity *in vivo*. **A**. Schematic of the experimental set up of the genetic screen in the *Drosophila* FUS model. Candidate genetic modifiers were identified in an unbiased screen, using the BDSC deficiency kit, the Exelixis kit, as well as specific RNAi lines and null mutants against the genes of interest. A complimentary literature-based approach was also used. Eclosion of a fly from the pupal case was categorized as a rescue. In total, 24 genes were identified as candidate modifiers of FUS toxicity. **B**. RNAi-mediated genetic knockdown of *sgg* rescues the FUS-induced fly eclosion phenotype. *W+v-w1118* crossed to FUS serves as a control. (N=10 crosses/condition) **C**. RNAi-mediated genetic knockdown of *mts* rescues the FUS-induced fly eclosion phenotype. *w1118* crossed to FUS serves as a control. (N=10 crosses/condition) **D**. Pharmacological inhibition of GSK3 by lithium chloride (LiCl) rescues the eclosion phenotype in FUS flies. D42-Gal4/+ serves as a control. (N=5 crosses/condition) **E**. Pharmacological inhibition of PP2A by okadaic acid (OA) rescues the eclosion phenotype in FUS flies. D42-Gal4/+ serves as a control. (N=6 crosses/condition). Values in B, C, D and E are represented as the mean ± SEM. Statistical comparisons between controls and RNAi conditions (B, C) were determined using one-way ANOVA with Sidak’s multiple comparisons test or Kruskal-Wallis test with Dunn’s multiple comparisons for each group. Statistical comparisons between controls and treated conditions (D, E) were determined using one-way ANOVA with Sidak’s multiple comparisons or Kruskal-Wallis test with Dunn’s multiple comparisons. (*p<0.05, **p<0.01, ***p<0.001, ****p<0.0001, ns = not significant)

In order to identify which genes are mediating the rescue effect of the large deficiencies, we first screened with smaller deletions. In total, we screened 167 smaller deletions and found 24 that modified the pupal lethality. In order to identify the actual genes responsible for the modifying effects, we used RNAi lines from the ‘VIENNA *Drosophila* research center’ (VDRC) as well as lines carrying mutations in candidate genes which were likely to result in loss of function. We found 18 modifying genes out of 88 extensively tested RNAi lines that attenuated mutant FUS-induced pupal lethality (Table S1). Among these genes were *mts*, the *Drosophila* ortholog of human *PP2A-C*, and *sgg*, the *Drosophila* ortholog of human *GSK3β*.

As a complimentary approach to identifying modifying genes, we searched the available literature and found 24 candidate genes which have previously been linked to FUS-associated ALS. Screening this list using RNAi lines and putative null alleles allowed us to identify 6 more genes involved in FUS induced neurotoxicity, among them *sgg* (Table S1). Thus, we ended up with a total of 24 candidate modifiers, with *mts* being identified through the unbiased approach, and *sgg* through both approaches.

### Genetic and pharmacological inhibition of mts and sgg rescue eclosion and lifespan in flies

Our approach yielded a total of 24 genes which modify FUS toxicity when their expression is reduced. To prioritize hits for follow up, we investigated whether any of these genes fall into a common pathway. Mts is the catalytic subunit of the PP2A phosphatase complex in *Drosophila*, whilst sgg is the ortholog of GSK3β and has recently been linked to ALS/FTD^15,16^. PP2A has been proposed to directly reduce GSK3β inhibitory phosphorylation in human cells, and thus act as an activator of GSK3β^17^.

As *mts* and *sgg* were candidate modifiers of FUS-toxicity, we sought to confirm their modifying action in the FUS flies. We first backcrossed several RNAi lines against sgg and mts into a suitable control genetic background (see methods) for 6 generations to avoid genetic background effects. Wild-type (WT) FUS and mutant R521G FUS were expressed in fly MNs using the D42-Gal4 driver along with *mts* or *sgg* RNAi and eclosion was scored. Knockdown of *sgg* led to a rescue of the FUS fly eclosion phenotype for both WT and R521G FUS-expressing flies for all 4 RNAi lines tested (Fig. 1B). Knockdown of *mts* led to a rescue for eclosion in WT and R521G FUS-expressing flies for 2 out of 3 of the tested RNAi lines (Fig. 1C). We next tested the consequence of feeding pharmacological inhibitors to the flies. For this we used lithium chloride (LiCl), a well-known and widely used GSK3 inhibitor^18,19^, and okadaic acid (OA), a specific inhibitor of PP2A in human cells at concentrations of 1-10nM^20,21^. Addition of LiCl to the fly food led to a higher percentage of eclosion for flies expressing FUS in their MNs between a range of doses of 5-15mM LiCl (Fig. 1D). LiCl only affected eclosion of driver-only (D42-Gal4/+) control flies above 15mM (Fig. 1D), before becoming developmentally lethal at 100mM (not shown), suggesting that it is well tolerated by the model. OA similarly rescued the eclosion phenotype of FUS flies, with the best rescue occurring between approximately 10 and 25nM in fly food (Fig. 1E). Toxicity of OA was only observed at 50nM and above (Fig. 1E).

To further validate the modifying ability of *mts* and *sgg*, we tested the effect of their knockdown on the lifespan of FUS-expressing flies. We expressed FUS WT or R521G in adult MNs using the D42-Gal4 driver, but to avoid developmental lethality, we included a temperature sensitive Gal80 allele (TS-Gal80), and allowed the flies to develop at 18°C where FUS expression was suppressed, before moving adults to 25°C to induce expression. We have previously shown that expression of either WT or mutant R521G FUS in this manner, significantly reduces the lifespan of the flies compared to healthy controls, leading to death with a median lifespan of approximately 3 weeks^9^. We found that RNAi-mediated knockdown of *mts* or *sgg* led to an extension of the FUS fly lifespan with all of the RNAi lines tested (Fig. 2A, B). Importantly, knockdown of *sgg* and *mts* did not alter the levels of FUS protein in the heads of the flies used for lifespan (Fig. 2C, D), suggesting that they exert their effects independently of FUS expression or stability.

**Fig. 2.**
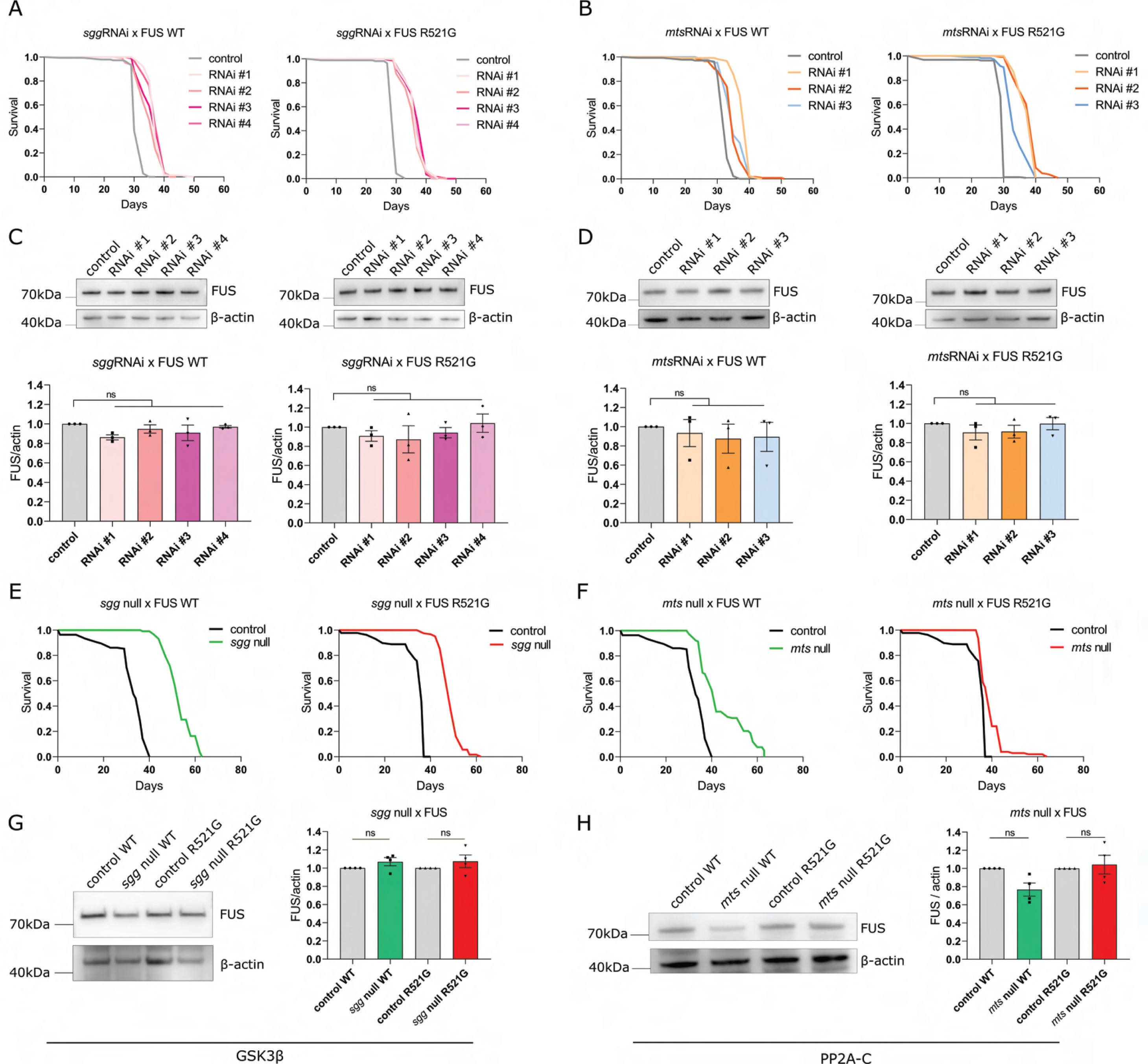
Inhibition of *sgg* or *mts* extends the lifespan of FUS flies. **A**. RNAi-mediated genetic knockdown of *sgg* extends the shortened lifespan of FUS WT and R521G *Drosophila* at 25℃. *W+v-w1118* crossed to FUS serves as control. (log-rank test, see Table S2 for statistical information) **B**. RNAi-mediated genetic knockdown of *mts* extends the shortened lifespan of FUS WT and R521G *Drosophila* at 25℃. *w1118* crossed to FUS serves as control. (log-rank test, see Table S2 for statistical information) **C, D**. Western blotting and quantification show that FUS expression remains unaltered in *Drosophila* heads after RNAi-mediated knockdown of *sgg* (C) or *mts* (D) (N=3, mean ± SEM, one-way ANOVA). **E**. A heterozygous null mutation in *sgg* (sgg^1^, Bloomington 4095) leads to a pronounced extension of the shortened lifespan of FUS WT and R521G flies at 25℃. *w1118* crossed to FUS serves as control. (log-rank test, see Table S3 for statistical information) **F**. A heterozygous null mutation in *mts* (^mtsXE2258^, Bloomington 5684) leads to a pronounced extension of the shortened lifespan of FUS WT and R521G flies at 25℃. *w1118* crossed to FUS serves as control. (log-rank test, see Table S3 for statistical information) **G, H**. Western blotting and quantification show that FUS expression remains unaltered in FUS WT and R521G *Drosophila* heads after heterozygous knockout of *sgg* (G) or *mt*s (H) (N=4, mean ± SEM, one-way ANOVA). *p<0.05, **p<0.01, ***p<0.001, ****p<0.0001, ns = not significant

As an alternative approach to investigate whether a stronger reduction of *mts* or *sgg* would lead to a more pronounced extension of the lifespan, we used null mutant lines of the two genes. As homozygous loss of function of these genes is developmentally lethal, we introduced a heterozygous loss of function. We observed a pronounced lifespan extension after heterozygous loss-of-function of *sgg* and *mts*, confirming the findings using RNAi (Fig. 2E, F). We observed an overall unaltered abundance of FUS protein after *sgg* and *mts* heterozygous knockout (Fig. 2G, H), and we additionally confirmed an approximate 50% reduction of sgg protein levels in the heterozygous mutant flies (Fig. S1). As antibodies against mts are not available, it was not possible to confirm the level of mts protein in the mts null mutant line.

Altogether, our results show that genetic and pharmacological inhibition of either mts or sgg can rescue FUS-induced developmental neurotoxicity. Moreover, genetic reduction of mts or sgg activity in adult flies can extend lifespan. These two modifying genes appear to act independently of FUS expression. We next sought to determine whether inhibition of the human orthologs of these genes can rescue toxicity in a human cellular model.

### Pharmacological inhibition of PP2A and GSK3 rescue hallmark ALS-associated phenotypes in iPSC-derived sMNs

To further confirm the modifying capacity of PP2A and GSK3, we investigated whether their inhibition could rescue hallmark FUS-ALS phenotypes. Cytoplasmic mislocalization of FUS is a pathological hallmark of ALS that has been linked with protein toxicity and neuronal death^6,22,23^. We therefore decided to investigate whether pharmacological inhibition of PP2A and GSK3 could rescue this phenotype. To assess the modifying effect of PP2A and GSK3 on mislocalization, we used a well-established iPSC-line with a *de novo* point mutation (P525L) in *FUS* from a 17-year old ALS patient^4^. This patient line was systematically compared with its corresponding CRISPR-Cas9 gene-edited isogenic P525P control^3,4^. In P525P FUS isogenic controls, FUS was mostly localized in the nucleus (Fig. 3A). In P525L FUS mutant iPSC-derived sMNs we observed cytoplasmic mislocalization of FUS, which was rescued after a 48h treatment with 1mM LiCl (Fig. 3B). Given that LiCl can produce off-target effects^19^, we also validated our findings using a highly selective GSK3 inhibitor, called tideglusib or NP12^24–27^. Tideglusib (TD) is a non-ATP competitive inhibitor of GSK3, which was well tolerated in phase 2 clinical trials for progressive supranuclear palsy (PSP)^28^ and is investigated in clinical trials for treating GSK3 hyperactivity in Alzheimer’s disease^25,26^. Recently, it has also been suggested as a potential treatment for ALS^15,16,24,29^. Excitingly, a 48h treatment of the P525L FUS MNs with 15μM tideglusib led to a significant rescue of FUS mislocalization too (Fig. 3B), confirming that the phenotype alleviation is not due to aspecific effects of LiCl, but thanks to GSK3 inhibition. To inhibit PP2A, we treated the FUS MNs with 1nM okadaic acid (OA) for 72h, resulting in a significant rescue of FUS mislocalization (Fig. 3B).

**Fig. 3.**
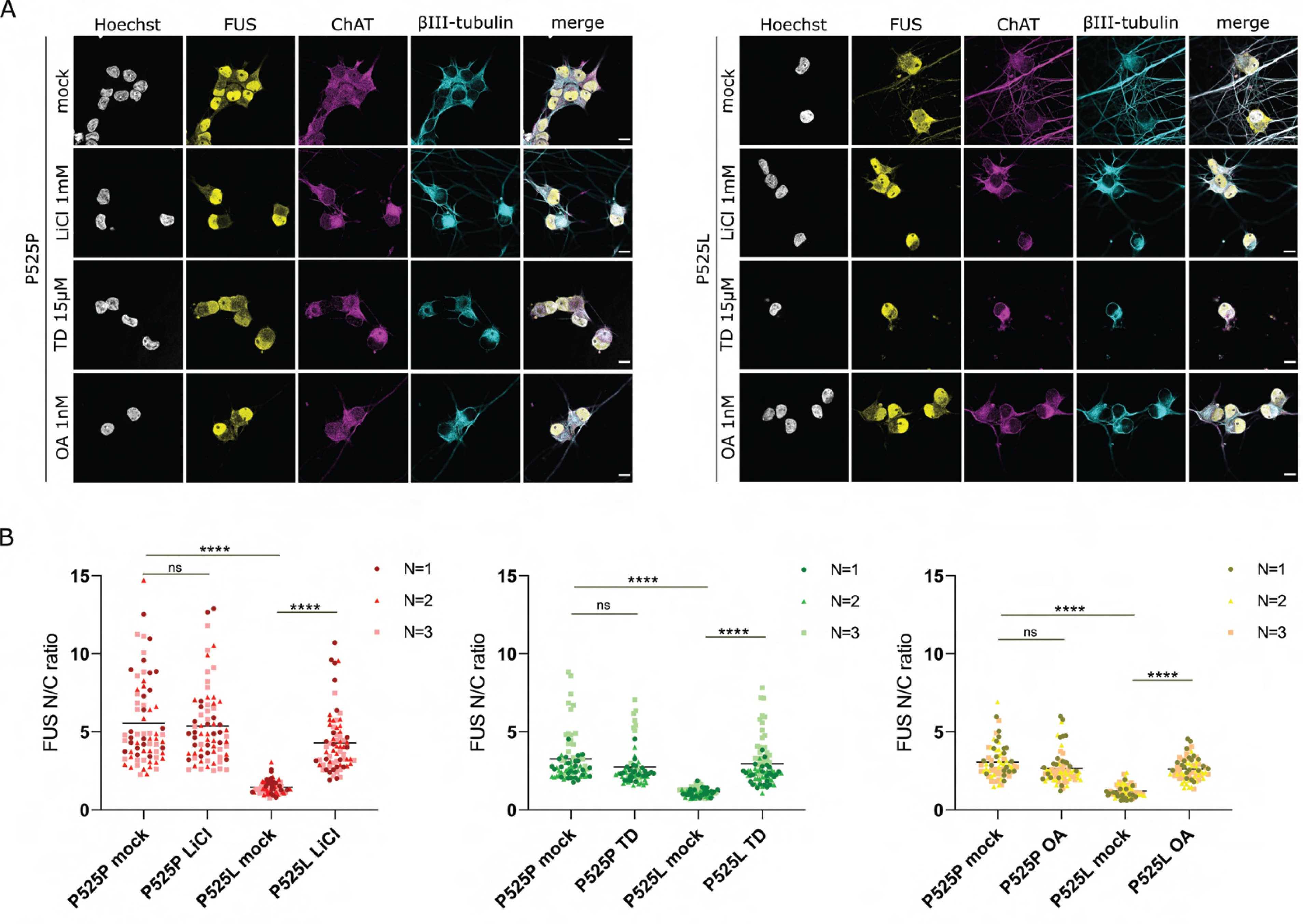
PP2A and GSK3 pharmacological inhibition rescue FUS cytoplasmic mislocalization in patient iPSC-derived motor neurons. **A**. Representative confocal images showing FUS distribution in patient iPSC-derived motor neurons with the P525L mutation, as well as in P525P isogenic controls after treatment with lithium chloride (LiCl, 1mM 48h), tideglusib (TD, 15μM 48h) or okadaic acid (OA, 1nM 72h). Scale bars = 10μm. **B**. Quantification of nuclear/cytoplasmic ratios (N/C ratio) fluorescent intensity of FUS in motor neurons of panel A shows a rescue of FUS mislocalization after treatments. Each dot represents one analyzed cell. Three different colors indicate data combined from three independent differentiations (N=70-80 cells). Data are shown as Grand mean; Kruskal-Wallis with Dunn’s multiple comparisons test. ****p<0.0001, ns = not significant.

We next determined the effect of pharmacologically inhibiting PP2A and GSK3 on NMJ formation. For this, we used a human-derived co-culture system that is well-established in our lab, combining iPSC-derived sMNs and myotubes in microfluidic devices, which allow us to study a functional human NMJ in a compartmentalized system^4,30,31^ Mutant FUS-associated NMJ impairment is a phenotype well-characterized in this model, as we observe a lower number of NMJs per myotube in the P525L mutant when compared to the P525P isogenic control^4^. On day18 of differentiation, we treated our motor neuron – myotube co-culture system with LiCl (1mM 48h), TD (15μM 48h) or OA (1nM 72h) in both the motor neuron and myotube compartments. On day28 we fixed the cells and assessed the number of NMJs by performing immunocytochemistry against NMJ markers. Using confocal microscopy, we observed that all three treatments had a positive effect on NMJ formation in the P525L mutant co-cultures, demonstrating the strong modifying capacity of GSK3 and PP2A (Fig. 4).

**Fig. 4.**
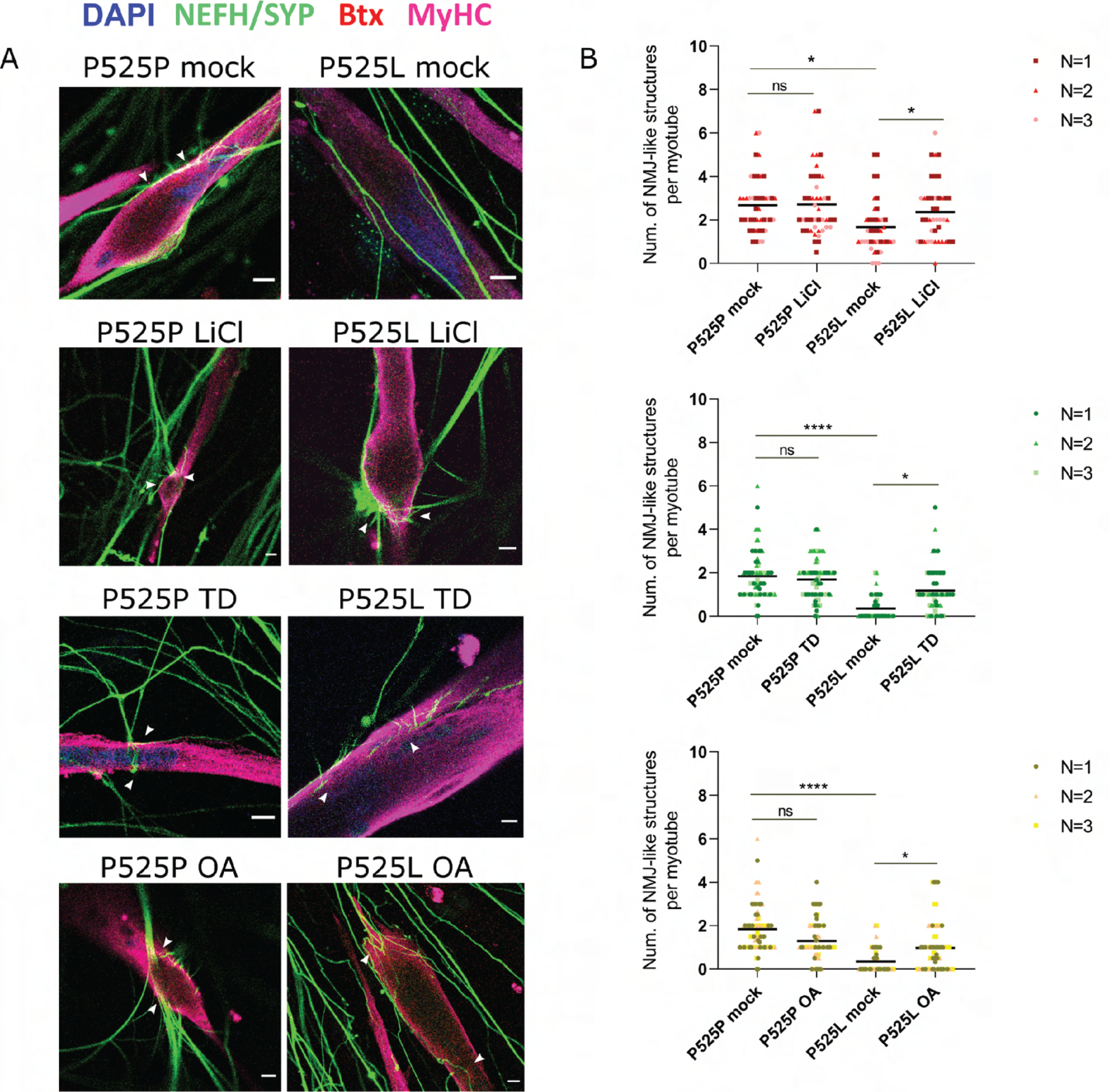
PP2A and GSK3 inhibition improve ALS-associated NMJ impairments. **A.** Representative confocal micrographs of NMJs (indicated with arrowheads) with LiCl, TD and OA treatments. NMJs are impaired in FUS-P525L ALS, and inhibition of PP2A or GSK3 improves the phenotype. Scale bars = 10μm **B**. Quantification of NMJ-like structures for all conditions, based on the neurite/presynaptic marker morphology and/or based on Btx (red)-SYP/NEFH (green) co-localization per myotube. Data are shown as Grand mean; N=3 biological replicates; Kruskal-Wallis test with Dunn’s multiple comparisons test. *p<0.05, ****p<0.0001, ns = not significant

Due to the polarization and length of sMNs, a proper regulation of axonal transport is essential for their function^32^. Defects in axonal transport are considered an early event in ALS pathogenesis, preceding axon retraction and denervation of the lower motor neuron or muscle^33,34^. We investigated whether PP2A or GSK3 inhibition could rescue mitochondrial transport deficits in our P525L FUS iPSC-derived sMNs. We performed live cell imaging of mitochondria at day30 of differentiation. We quantified the total number of mitochondria per 100μm of neurite, as well as the total number of stationary and moving mitochondria (Fig. S2), allowing us to calculate the percentage of moving mitochondria. Compared to P525P FUS isogenic controls, the percentage of moving mitochondria was significantly lower in P525L sMNs (Fig. 5). After treating the cells with LiCl for 48h, to inhibit GSK3, tracking analysis showed a clear rescue of mitochondrial transport defects (Fig. 5A, D). Moreover, a similar rescue of transport was observed after treatment with tideglusib (Fig. 5B, E), confirming the modifying capacity of GSK3 inhibition. In addition, OA treatment for 72h similarly led to a significant improvement of mitochondrial transport (Fig. 5C, F). Although we occasionally observed small differences in the total number of mitochondria per neurite, the drug treatments consistently reduced the number of stationary mitochondria and increased the number of motile mitochondria (Fig. S2). Altogether, our results demonstrate that PP2A and GSK3 are modifiers of FUS-induced neurotoxicity both in *Drosophila* and human cellular models.

**Fig. 5.**
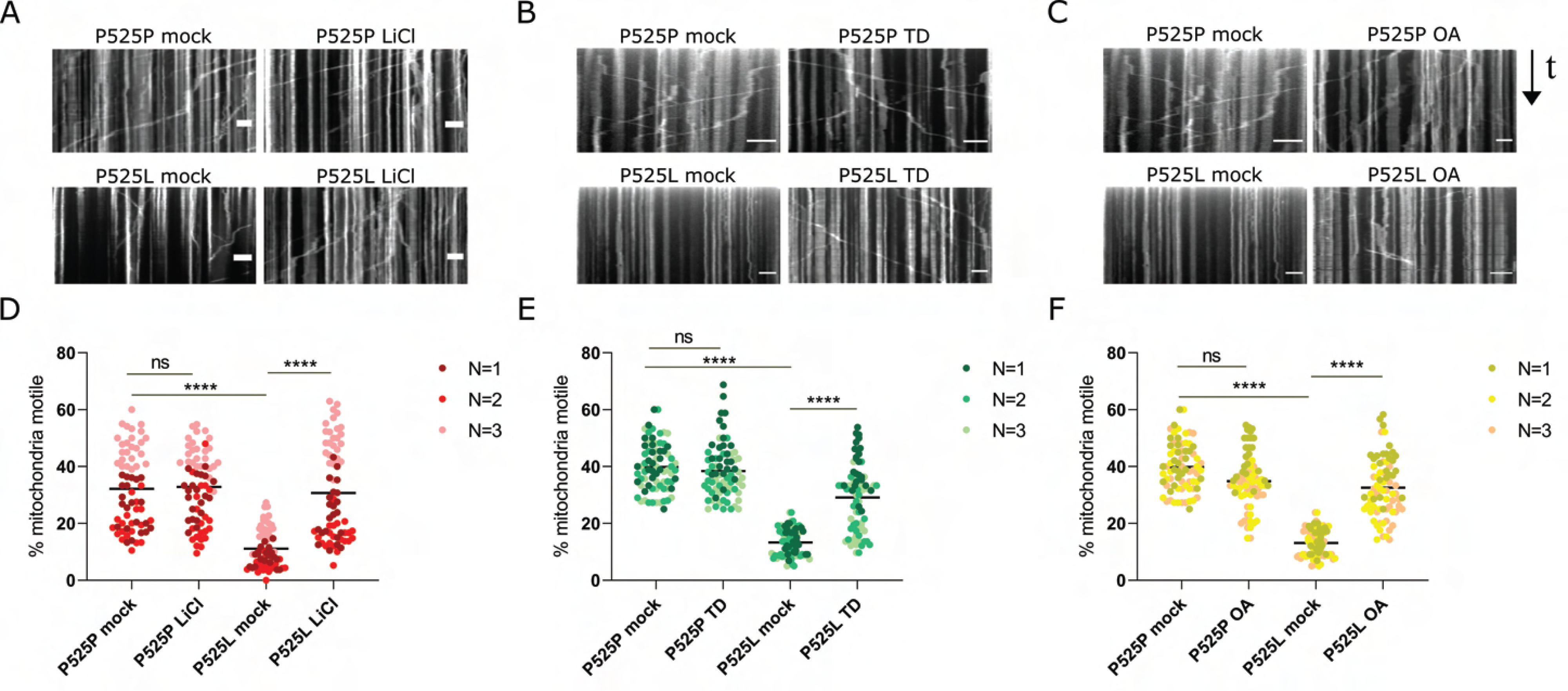
Pharmacological inhibition of PP2A and GSK3 rescue mitochondrial transport deficits. **A-C.** Example kymographs (time-distance plots) of mitochondria (MitoTracker Green) after treatment of 30-day old P525P isogenic control motor neurons and P525L mutant motor neurons with 1mM lithium chloride (LiCl) for 48h (A), 15μM tideglusib (TD) for 48h (B) and 1nM okadaic acid (OA) for 72h (C). Scale bars 30μm **D-F.** Percentage of mitochondria that are motile in the motor neurons (day30) comparing isogenic control and mutant with or without LiCl (D), TD (E) and OA (F). Data shown as Grand Mean; N=3; Kruskal-Wallis with Dunn’s multiple comparisons test. ****p<0.0001, ns = not significant

### GSK3 is hyperactive in FUS-ALS due to reduced inhibitory phosphorylation

Recently, reduced GSK3β inhibitory phosphorylation was observed in a FUS mouse model^15^, suggesting that GSK3β may become hyperactive in response to dysfunctional FUS protein. To further investigate GSK3 hyperactivity in disease, we started with our FUS fly model. To determine whether GSK3 is hyperactive in *Drosophila*, we used nSyb-Gal4 combined with Gal80-ts to drive FUS expression pan-neuronally in adult flies for 7 days, and assessed sgg expression and phosphorylation by western blotting (Fig. 6). Consistent with previous reports^35^, we observed two major bands of sgg protein in *Drosophila* head lysates corresponding to isoforms SGG10 and SGG39 of the protein. Upon WT and R521G FUS expression, we noticed a significant reduction in the levels of phospho-sgg, while the levels of total-sgg protein remained unaltered (Fig. 6A, B). The reduced inhibitory phosphorylation therefore suggests that GSK3 is hyperactive in our FUS flies. To translate our *Drosophila* findings to human cells, we performed western blotting for phospho-GSK3α/β at residue serine 21/9 in our patient iPSC-derived FUS sMNs on day 30 of differentiation. Consistently to what we observed in the fly model, we found that phospho-GSK3α/β was significantly reduced in mutant P525L FUS MNs compared to the P525P isogenic controls (Fig. 6C, D), demonstrating that GSK3 inhibitory phosphorylation is also reduced in FUS patient sMNs and suggesting that GSK3 is hyperactive in this model as well.

**Fig. 6.**
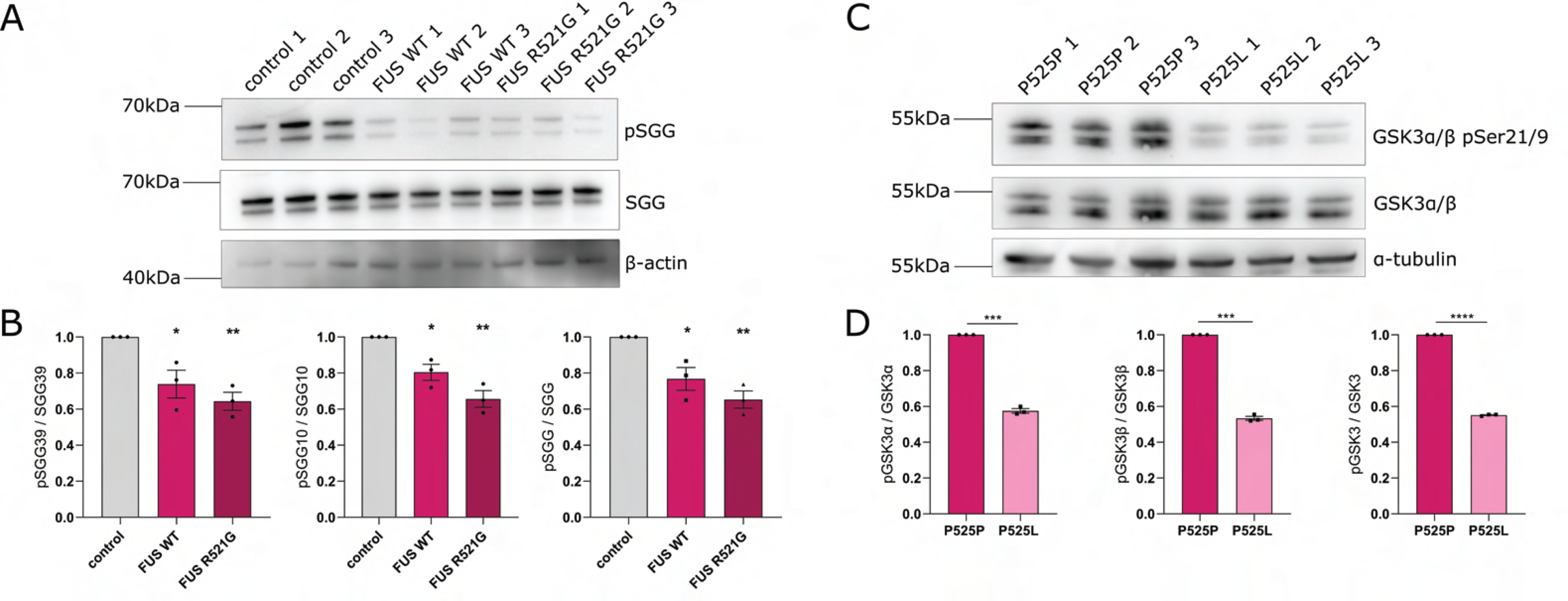
GSK3 is hyperactive in FUS-ALS due to reduced inhibitory phosphorylation. **A**. Western blot showing phospho-sgg and total-sgg protein in control vs FUS WT and R521G flies. Control was *w1118* crossed to nSyb-Gal4. In *Drosophila*, the two splice isoforms of GSK3-β, SGG39 (upper band) and SGG10 (lower band), are seen. Beta-actin serves as a loading control. **B**. Quantification of panel A shows that phosphorylation of SGG is reduced in FUS WT and R521G *Drosophila*, for the total-SGG protein and for the individual SGG39 and SGG10 isoforms, while the levels of SGG protein remain unaltered overall. (N=3, mean ± SEM, one-way ANOVA with Sidak’s multiple comparisons test) **C**. Western blot for GSK3α/β pSer21/9 in iPSC-derived motor neurons from FUS-P525L patients and in their CRISPR-corrected P525P isogenic controls. Ser21/9 inhibitory phosphorylation of GSK3α/β appears reduced in FUS P525L patient motor neurons compared to isogenic controls, while the total levels of GSK3α/β remain unaltered among the conditions. The numbers 1, 2 and 3 represent three independent differentiations. Alpha-tubulin serves as a loading control. **D**. Quantification of the western blot of panel C shows reduced Ser21 and Ser9 inhibitory phosphorylation for GSK3 α and β respectively. (N=3, mean ± SEM, two-tailed paired t-test) *p<0.05, **p<0.01, ***p<0.001, ****p<0.0001

### PP2A acts upstream of GSK3 affecting its phosphorylation, and GSK3 hyperactivity alone is insufficient to drive toxicity

PP2A has been suggested to modify the phosphorylation of GSK3, affecting its activity^17^. To assess a possible PP2A-GSK3 interaction in our model, we first tested whether PP2A can regulate GSK3 phosphorylation in *Drosophila*. To determine whether mts controls sgg phosphorylation, we overexpressed mts using the pan-neuronal nSyb-Gal4 driver. After aging the progeny for 7 days, we assessed GSK3 phosphorylation using western blotting. We found that *mts* overexpression led to significantly decreased levels of inhibitory phosphorylation of both major sgg isoforms, suggesting that PP2A acts upstream of GSK3 in flies and causes its dephosphorylation (Fig. 7A, B). To determine whether GSK3 hyperactivity is sufficient to lead to neurodegeneration, we overexpressed either wild-type sgg (*UAS-sgg*) or a constitutively active mutant in which the serine 9 phosphorylation site had been replaced with alanine (*UAS-sgg S9→A*). Overexpression of these sgg variants in fly MNs, using the D42-Gal4 driver, caused an eclosion phenotype, milder than the one observed after FUS expression, suggesting that GSK3 hyperactivity alone can drive toxicity but may not be fully responsible for the strong eclosion defect that we observed in FUS-expressing flies (Fig. 7C). Recapitulating the rescue that we previously saw, addition of LiCl in the food rescued the mild eclosion defect induced by sgg overexpression, increasing eclosion rates from ∼60% to ∼85% (Fig. 7C). On the contrary, LiCl had no effect on the eclosion of the flies expressing the constitutively active form of sgg, suggesting that lithium inhibits sgg via increasing its phosphorylation at Ser9 (Fig. 7C). To determine whether PP2A inhibition could rescue sgg hyperactivity, we added OA to the food of sgg-overexpressing flies. OA significantly increased eclosion of *UAS-sgg* flies, but had no effect on the eclosion of the *UAS-sggS9→A* flies (Fig. 7D). These data indicate that PP2A and GSK3 act in a common pathway, with PP2A lying upstream of GSK3. Even more importantly, we conclude that PP2A can alter GSK3 inhibitory phosphorylation on this specific Serine9 residue.

**Fig. 7.**
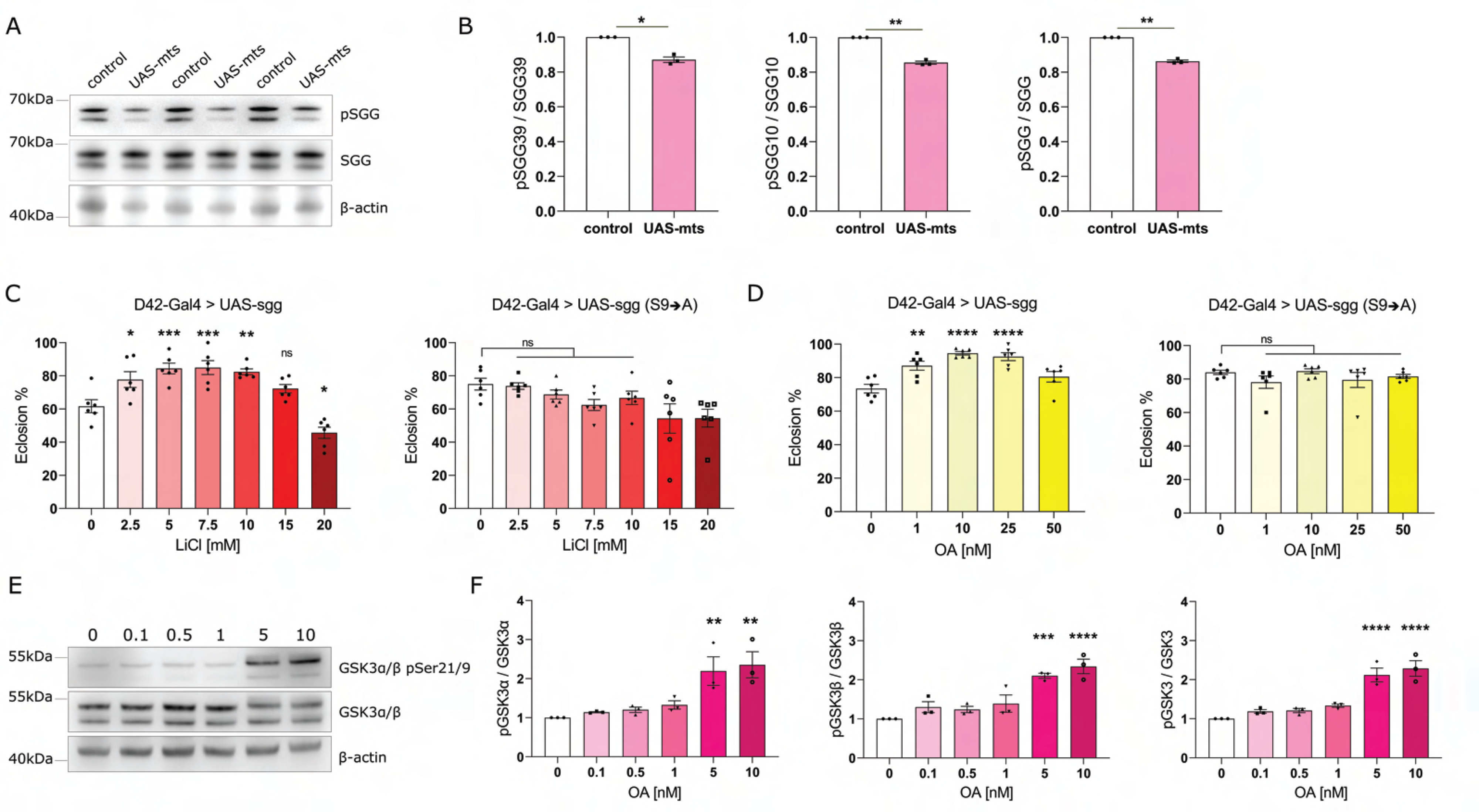
PP2A acts upstream of GSK3, affecting its inhibitory phosphorylation. **A**. Western blot for phospho-sgg and total-sgg protein in control vs flies overexpressing mts pan-neuronally. Control was *w1118* crossed to nSyb-Gal4. In *Drosophila*, the two splice isoforms of GSK3-β, SGG39 (upper band) and SGG10 (lower band), are seen. Beta-actin serves as loading control. **B**. Quantification of panel A shows that phosphorylation of SGG is reduced after mts overexpression in the *Drosophila* brain, for the total-SGG protein and for the individual SGG39 and SGG10 isoforms, while the levels of SGG protein remain unaltered overall. (N=3, mean ± SEM, two-tailed paired t-test). **C**. Overexpression of sgg in the fly motor neurons causes a mild eclosion defect, with ∼60% flies eclosing. LiCl in the fly food rescues the eclosion deficit to ∼85-90%. Overexpression of a constitutively active form of sgg (S9◊A) leads to a mild eclosion defect (∼75-80%) and LiCl has no effect on this deficit. Statistical comparisons between control (0) and treated conditions (2.5-20) were determined using one-way ANOVA with Sidak’s multiple comparisons (N=6 vials/condition). **D**. Overexpression of sgg in the fly motor neurons causes a mild eclosion defect. OA in the fly food rescues the eclosion deficit to ∼90-95%. Overexpression of a constitutively active form of sgg (S9◊A) also leads to a mild eclosion defect (∼80%) and OA has no effect on this deficit. Statistical comparisons between control (0) and treated conditions (1-50) were determined using one-way ANOVA with Sidak’s multiple comparisons or Kruskal-Wallis with Dunn’s multiple comparisons test (N=6 vials/condition). **E**. Western blot for GSK3α/β pSer21/9 in SH-SY5Y cells treated with OA 0-10nM. Ser21/9 inhibitory phosphorylation of GSK3α/β starts to increase with an increasing dose of OA, while the total levels of GSK3α/β remain generally unaltered. Beta-actin serves as the loading control. **F**. Quantification of the western blot of panel E shows a significant increase of Ser21/9 inhibitory phosphorylation for GSK3α/β after treatment with 5nM OA and 10nM OA. (N=3, mean ± SEM, one-way ANOVA with Sidak’s multiple comparisons) *p<0.05, **p<0.01, ***p<0.001, ****p<0.0001, ns = not significant

These results suggest that in *Drosophila*, one role of PP2A is to modify the activation state of GSK3. To test whether this interaction is conserved in human cells, we treated SH-SY5Y cells with OA, in a concentration range of 0-10nM. With increasing inhibition of PP2A, we observed increasing levels of phospho-GSK3α/β, indicating that PP2A controls GSK3 phosphorylation in flies as well as in human cells (Fig. 7E, F).

### Increased expression of kinesin-1 is sufficient to rescue FUS toxicity in *Drosophila* models

Given that we observe defects in the trafficking of mitochondria in iPSC-derived sMNs, we sought to assess whether increasing axonal transport in our *Drosophila* model might rescue toxicity. Kinesin-1 is strongly associated with neurodegeneration and ALS^36,37^. *Drosophila* has a single kinesin-1 heavy chain ortholog (*Khc*) and a single light chain ortholog (*Klc*). Loss of function of *Khc* and *Klc* results in motor dysfunction in *Drosophila* larvae, leading to axonal swellings packed with mitochondria and other fast axonal transport cargo, suggesting that these genes are essential for transport of organelles^38,39^. Previously, Vagnoni *et al*. demonstrated that introduction of a genomic rescue construct consisting of a P-element that carries a 7.5 kb genomic fragment including the 3.5 kb kinesin heavy chain transcription unit (P[Khc+]) into wild-type flies can rescue age-related axonal transport deficits^40^. Despite only encoding Khc, this rescue construct has also been reported to increase Klc at the protein level via an unknown mechanism^38^.

To rule out genetic background effects, we backcrossed P[Khc+] into the *w1118* control background for 6 generations and crossed it to the FUS WT and R521G eclosion stock. P[Khc+] rescued the eclosion phenotype of our FUS flies (Fig.8A).

**Fig. 8.**
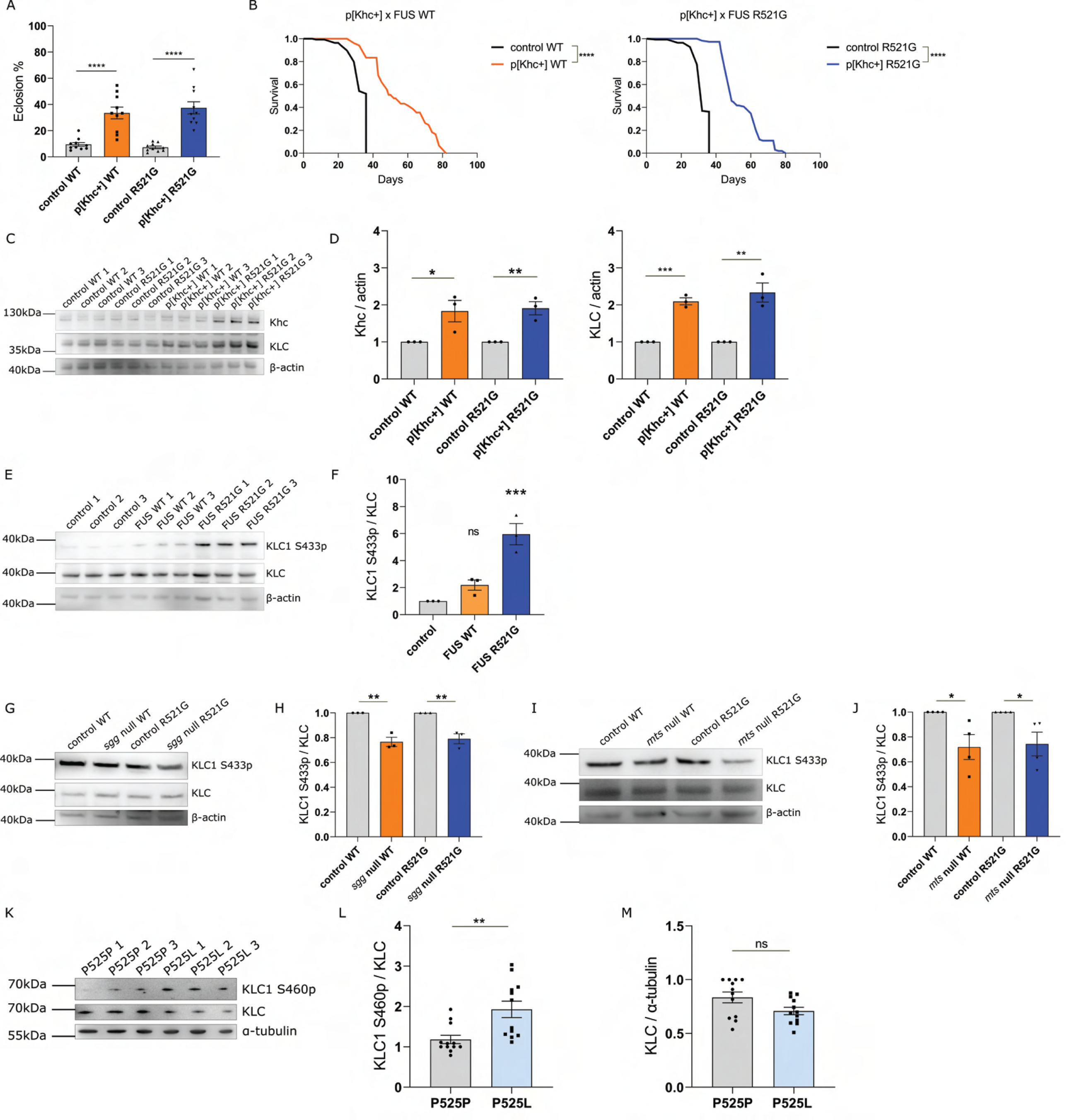
Kinesin appears dysfunctional in FUS-ALS, perhaps due to hyper-phosphorylation by GSK3. **A.** Overexpression of Khc using the p[Khc+] stock rescues the FUS-induced fly eclosion phenotype. *w1118* crossed to D42Gal4-driven FUS serves as control. (N=10 crosses/condition) **B**. Overexpression of Khc strongly extends the shortened lifespan of FUS WT and R521G *Drosophila* at 25℃. *w1118* crossed to FUS serves as control. (see Table S4 for statistical information) **C**. Western blot for Khc and Klc in FUS flies vs FUS flies with p[Khc+]. Beta-actin serves as loading control. **D**. Quantification of the western blots of panel C show a significant increase in Khc and Klc levels (N=3). **E**. Western blot of KLC1 S433 phosphorylation in control w1118 VS FUS-expressing flies. **F**. Quantification of the western blot of panel E shows increased levels of KLC1 S433p in FUS flies. **G**. Western blot of KLC1 S433p in FUS flies with sgg heterozygous knockout (N=3). **H**. Quantification of panel G shows that KLC1 S433p levels decrease after sgg inhibition. **I**. Western blotting assessing KLC1 S433p in FUS flies with mts heterozygous knockout (N=3). **J**. Quantification of panel I shows that KLC1 S433p levels decrease after mts inhibition. **K.** Western blotting in FUS P525P and P525L MNs, assessing KLC1 S460 phosphorylation. **L**. Quantification of panel K reveals increased KLC1 S460p in P525L patient motor neurons compared to isogenic controls (N=3 differentiations). **M.** Quantification from panel K shows a trend for reduced KLC1 levels in FUS P525L MNs. Values in A, D, H, J and L are represented as the mean ± SEM and statistical comparisons were determined using unpaired t-test. One-way ANOVA with Sidak’s multiple comparisons test was used for panel F. *p<0.05, **p<0.01, ***p<0.001, ****p<0.0001, ns = not significant

We also performed lifespan experiments and found that P[Khc+]-carrying FUS flies lived significantly longer than controls (Fig.8B). Subsequently, we performed western blotting on lysates from heads of flies used in the lifespan experiment and observed that both Khc and Klc levels were significantly increased by the introduction of the P[Khc+] rescue construct (Fig.8C, D).

These results strongly suggest that increasing the levels of kinesin light and heavy chain is beneficial in our fly model.

### Kinesin light chain phosphorylation is increased by FUS dysfunction and rescued by inhibition of PP2A/GSK3

Our results showed that inhibition of either GSK3 or PP2A can rescue FUS-induced toxicity in both flies and iPSC-derived sMNs. FUS is known to be phosphorylated by a number of DNA-damage associated kinases like ATM^41^. To test whether PP2A could be a phosphatase of FUS, we induced FUS phosphorylation by treating HEK293T cells for 2h with 10nM calicheamycin γ1 (CLM), an antibiotic that cleaves DNA and specifically induces DNA double strand breaks (DSBs)^42^. We observed robust FUS phosphorylation, visible as an increase in the apparent molecular weight of the protein after SDS-PAGE (Fig. S3A). To test whether PP2A could be affecting the phosphorylation of FUS, we treated the cells with 10nM OA directly after the induction of DNA damage and assessed whether OA treatment affected the dephosphorylation of FUS. We failed to observe any significant effect of OA on FUS dephosphorylation after DNA damage (Fig. S3C), suggesting that OA is acting independently of FUS phosphorylation. To investigate whether a direct physical contact exists between PP2A and FUS, regardless of phosphorylation, we performed pull down experiments. We transfected HEK293T cells with FLAG-tagged WT FUS and treated these with 10nM CLM for 2h. Although the pull down of the FLAG-tagged FUS was efficient, we were not able to detect PP2A on western blot analysis in normal or CLM treated conditions (Fig. S3B).

These results suggest that the modifying effect of PP2A does not depend on a direct physical interaction with FUS nor on its de-phosphorylating ability. This suggests that PP2A acts indirectly to induce FUS toxicity.

Given that PP2A/GSK3 inhibition appears unlikely to affect FUS phosphorylation directly, we wondered whether these proteins could be directly involved in some of the downstream pathologies that FUS induces.

Kinesin light chain is a substrate of GSK3β *in vivo*,^13,43,44^ suggesting that a potential GSK3 hyperactivity in FUS-ALS could lead to kinesin hyperphosphorylation and dysfunction/degradation. Recent studies showed that KLC1 serine-460 phosphorylation causes axonal transport deficits and contributes to neurodegeneration in Alzheimer’s disease^45^. Given that this phosphorylation site is well conserved in *Drosophila* (serine 433)^45^, we used western blotting to examine Klc S433 phosphorylation in FUS-expressing flies (Fig. 8E). After pan-neuronal expression of either WT or R521G mutant FUS, we observed that Klc S433 phosphorylation levels increased, particularly in R521G-expressing flies (Fig. 8F).

To test the modifying ability of PP2A and GSK3 on this downstream target, we used our *mts* and *sgg* null mutant lines and we examined the levels of Klc S433 phosphorylation after *mts* or *sgg* heterozygous knockout. Excitingly, we saw that the levels of Klc S433 phosphorylation decreased again after *mts* or *sgg* knockdown in both WT and mutant FUS flies (Fig. 8G, H, I, J). These data support our hypothesis that the changes in GSK3 phosphorylation lead to its hyperactivity, increasing kinesin-1 phosphorylation and culminating in dysfunction in FUS-ALS. Finally, to translate our *Drosophila* findings of FUS-associated kinesin dysfunction to humans, we performed western blotting in P525P isogenic and P525L mutant FUS iPSC-derived sMNs. In line with our fly data, we found that KLC1 serine-460 phosphorylation was increased in FUS mutant sMNs (Fig. 8K, L). Moreover, there was a tendency for KLC levels to be reduced in the mutant FUS sMNs on the blot, further indicating a possible dysfunction/degradation of KLC in FUS-ALS (Fig. 8K, M). Altogether, our data indicate a previously unknown link between GSK3 hyperactivity and kinesin dysfunction, leading to mitochondrial transport deficits in FUS-ALS.

## Discussion

The heterogeneity of ALS in terms of its pathology and clinical presentation complicates our understanding of the disease^46–49^. Since this variability may arise from the existence of disease modifying-genes, we performed a genome-wide screen in *Drosophila* to discover novel modifiers of FUS-ALS. We identified a total of 24 genes whose loss-of-function rescued the FUS eclosion defect. The genes *mts* (*PP2A-C*) and *sgg* (*GSK3β*) were two of those candidates which we pursued further. Genetic and pharmacological inhibition of *mts* or *sgg* rescued FUS-induced phenotypes in *Drosophila*, such as eclosion defects and reduced lifespan. Treatment of FUS iPSC-derived sMNs with PP2A or GSK3 inhibitors rescued hallmark ALS phenotypes, such as FUS cytoplasmic mislocalization, reduced NMJ formation and mitochondrial transport defects, further supporting our finding that PP2A and GSK3 are modifiers of FUS toxicity.

GSK3 is a ubiquitously expressed kinase, whose activity depends on its own inhibitory phosphorylation^50^. GSK3 hyperactivity has been linked to neurodegeneration before, mainly AD, and recent studies suggested a possible role of GSK3 in FUS and TDP-43 ALS^15,16,29^. Notably, loss of function of sgg has been shown to rescue neuronal dysfunction in a TDP-43 *Drosophila* model^16,51^. Moreover, our findings are in line with Choi *et al*., who recently suggested that sgg inhibition could rescue various phenotypes in *Drosophila* overexpressing human FUS^52^. PP2A is a multi-subunit phosphatase highly expressed in the brain, of which PP2A-C (mts) is the catalytic subunit. PP2A has been proposed to reverse GSK3 inhibitory phosphorylation in mammalian cells^53^, suggesting it may regulate GSK3 activity.

In both our *Drosophila* model as well as in iPSC-derived sMNs of FUS mutation carriers, we found reduced GSK3 phosphorylation. Recently, reduced inhibitory phosphorylation of GSK3 has also been demonstrated in a FUS mouse model^15^. These results suggest that FUS may directly regulate the level of GSK3 phosphorylation via an as-yet-unknown mechanism. Excitingly, lymphoblasts from sporadic ALS patients have also been shown to display reduced GSK3 phosphorylation^24^, implicating GSK3 hyperactivity in a broader patient population.

The relationship between PP2A and GSK3 is not well studied. Recent findings suggest that these two enzymes can affect the activities of each other, with PP2A affecting GSK3 phosphorylation^54^. In our study, we have developed multiple lines of evidence that PP2A affects the level of GSK3 inhibitory phosphorylation. In *Drosophila*, overexpression of the PP2A-C ortholog mts reduced GSK3 serine 9 phosphorylation, while okadaic acid, a PP2A inhibitor, rescued sgg overexpression-induced toxicity in a manner dependent on GSK3 phosphorylation at serine 9. In SHSY-5Y cells we observed a dose-dependent increase in GSK3α/β phosphorylation in response to okadaic acid. Whilst it is possible that the okadaic acid is also affecting PP1, a known GSK3 phosphatase^55,56^, okadaic acid has a higher specificity for PP2A over PP1 at these concentrations^57,58^, and the direct effect of mts overexpression suggests that PP2A is indeed directly involved in regulating the phosphorylation of GSK3.

Healthy mitochondrial transport is crucial for neuronal survival and homeostasis. Mitochondrial transport deficits are linked with several neurodegenerative disorders, such as Alzheimer’s disease, Huntington’s disease^59,60^, hereditary spastic paraplegia^61^, as well as ALS with *SOD1, TARDBP* and *FUS* mutations, and *C9orf72* with hexanucleotide repeats^7,8,12,62^. Axonal transport is mediated by the motor proteins kinesin and dynein^11^. The kinesin superfamily, mediating anterograde axonal transport, is encoded by 45 mammalian genes, with 38 of them being expressed in the nervous system^11,12^. We focused on kinesin-1, since the strongest genetic evidence connecting impaired axonal transport with neurodegeneration comes from mutations in the kinesin genes^36,37^. Kinesin-1 is formed from a dimer of two heavy chains (KHCs), encoded by KIF5A, KIF5B or KIF5C, which act as the motor domain of the protein, and a dimer of KLCs, which acts as an adaptor for cargoes^11,12,38^. KIF5A mutations were recently linked to a number of ALS cases, indicating that alterations in anterograde transport may be involved in the pathogenesis of ALS^37,63^. Moreover, swellings of organelles and proteins that specifically ensnare kinesin were found in the motor axons of ALS patients *post-mortem*^64,65^, and differential expression of kinesin isoforms was observed in the motor cortex of sALS patients^66^. Moreover, it was shown that loss of Klc or Khc function lead to disruption of axonal transport in *Drosophila* and similar axonal transport defects^38^.

Various kinases can phosphorylate motor proteins responsible for axonal transport, with a previous study suggesting that GSK3β can modulate kinesin-1-based transport by phosphorylating KLC2^67,68^. Inhibiting GSK3β activity in primary cortical neurons of a transgenic mouse model of AD led to restoration of Aβ-induced transport deficits^69^. We found that the reduced neuritic transport of mitochondria in P525L FUS iPSC-derived sMNs can be rescued by pharmacological inhibition of GSK3, either by LiCl or tideglusib. Moreover, PP2A inhibition by OA led to a similar rescue of the phenotype. These data suggest that PP2A and GSK3 could act as modifiers of FUS-induced toxicity by modulating mitochondrial transport. Khc/Klc overexpression in our FUS flies rescued both eclosion defects as well as shortened lifespan, suggesting a dysfunction of kinesin-mediated axonal transport in these flies. Using recently generated antibodies against KLC1 S460, a phosphorylation site which regulates KLC-cargo binding both in flies and humans^45,70^, we observed that FUS overexpression in *Drosophila* strongly increased KLC phosphorylation. We found a similar hyperphosphorylation in patient iPSC-derived sMNs, suggesting a conservation of mechanism. Previously, it has been suggested that KLC1 is phosphorylated at S460 by ERK in human cells^45^. Whilst the role of GSK3 has not been explored, the fact that sgg loss of function in *Drosophila* can regulate this event suggests that GSK3 is involved in S460 phosphorylation as well. It is somewhat surprising that mts loss-of-function reduced KLC S460p in *Drosophila*, given that okadaic acid is thought to increase kinesin-1 phosphorylation via ERK activation in human cells^45^. This difference may be because OA was applied by Mórotz *et al*. at 50nM, a dose sufficient to generally inhibit Serine/Threonine protein phosphatases, whilst our treatments were performed at 1nM. Our results point towards GSK3 hyperactivity and kinesin-1 dysfunction in FUS-ALS. Since GSK3 can phosphorylate kinesin-1, we concluded that GSK3 can modify axonal transport by acting on kinesin-1. GSK3 hyperactivity is induced after FUS expression, causing extensive kinesin-1 phosphorylation. This may lead to cargo release from the motor protein^68^ and may push kinesin into the proteasomal degradation pathway^13^, causing axonal transport defects (Fig. 9).

**Fig. 9.**
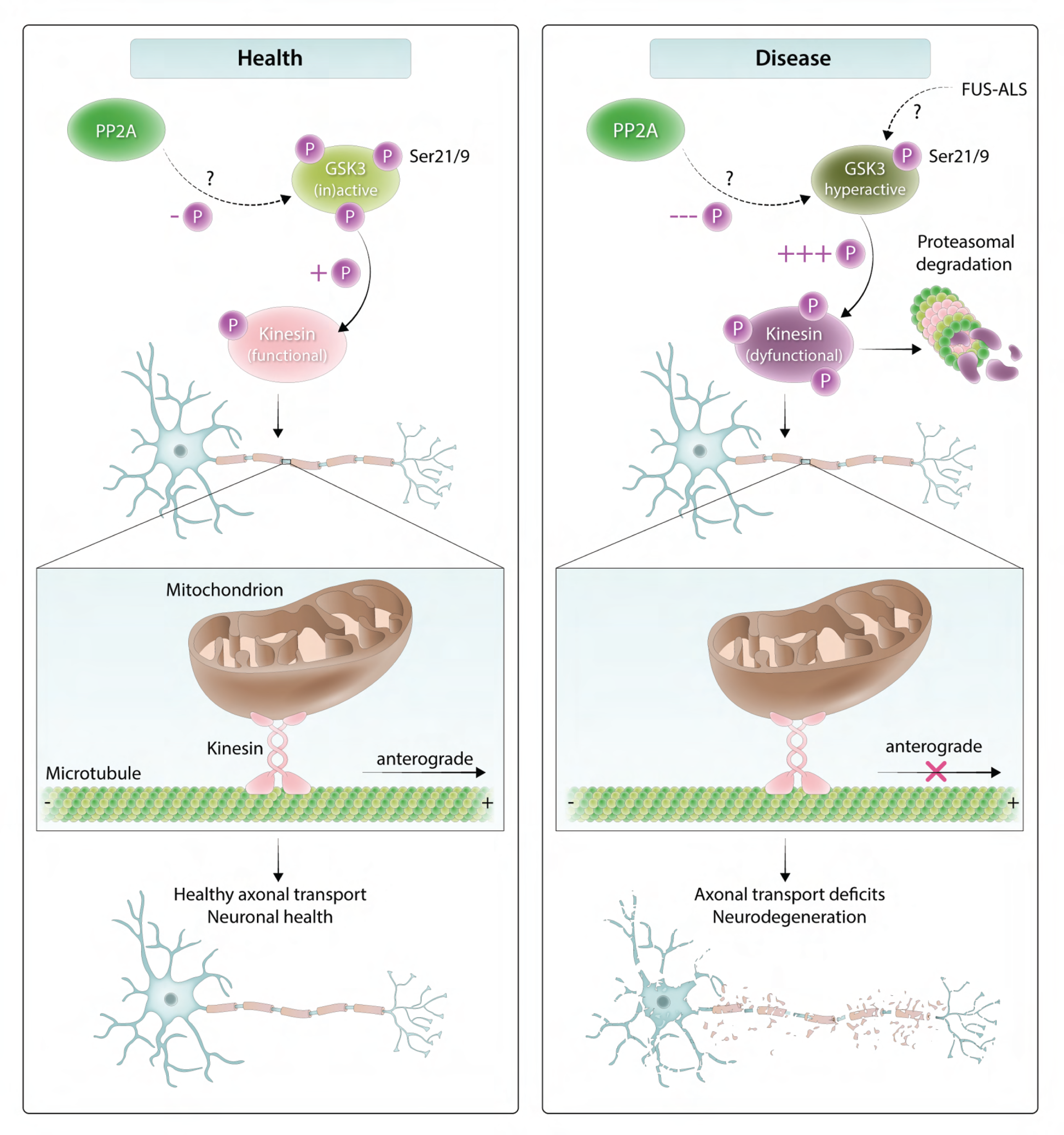
Working model for how GSK3 hyperactivity may be implicated in FUS-ALS. Our current data suggest that in a healthy cell, GSK3 inhibitory phosphorylation is at normal levels, properly regulating kinesin phosphorylation and allowing mediation of mitochondrial transport. However, in FUS-ALS, GSK3 becomes hyperactive, perhaps through the direct or indirect action of PP2A. Subsequently, GSK3 can lead to kinesin hyper-phosphorylation and cause cargo release or dysfunction/degradation, leading to mitochondrial transport deficits and neurodegeneration.

The proposed mechanism above still requires some points of clarification. Firstly, while KLC S460 phosphorylation is sufficient to prevent the interaction with some cargoes, it is currently unclear whether this phosphorylation event would be sufficient to drive the mitochondrial transport defect seen in our sMN cultures. It may be that other residues of KLC become hyper phosphorylated in the disease state affecting different aspects of cargo binding, and that S460p is simply a readout of a more global protein-wide hyperphosphorylation event. Further work performing unbiased mass spectrometry in our iPSC sMNs could be used to explore this further.

Secondly, while GSK3 hyperactivity has been implicated in multiple FUS-ALS models, we find in flies that overexpression of GSK3 (sgg) or its constitutively active form (S9→A) is insufficient to fully recapitulate the eclosion defect we observe in FUS-overexpressing flies. This may suggest that FUS overexpression induces multiple independent pathways leading to toxicity, including, but not limited to, GSK3 hyperphosphorylation. Further exploration of other modifying genes identified in our study may shed light on this.

We observed that GSK3 and PP2A inhibition was sufficient to return FUS to the nucleus in P525L mutant sMNs. This result may suggest that GSK3 inhibition is not working directly on axonal transport but rather on the localization of FUS itself. However, in contrast to this hypothesis, we and others have observed that removal of the nuclear localization sequence of human FUS is beneficial in flies^9,71^, suggesting that the rescue observed in *Drosophila* is not due to this mechanism. We speculate that the increased nuclear localization of FUS observed in the treated P525L sMNs may be due to a general increase in health of the cell encouraging FUS nuclear import. Future experiments which look at the effect of GSK3 and PP2A inhibitors on the localization of fluorescently tagged reporter proteins may help to determine whether these enzymes can affect nuclear import of FUS directly.

Finally, the use of GSK3 inhibitors, among them lithium^72,73^ and tideglusib^25^, is being investigated as a potential treatment for AD, in animal studies as well as in clinical trials^25,74,75^. We note that lithium carbonate has already been tested for the treatment of ALS patients, showing no clinical benefit. However, a genetic subgroup analysis indicated that lithium carbonate was beneficial in patients homozygous for the C allele of rs12608932 in UNC13A^76,77^, with a follow-up clinical trial recently announced^76^. Given the apparent heterogeneity in patient response to lithium, further research into its mode of action, and pre-clinical testing of other GSK3 inhibitors like tideglusib is warranted. In future it would be interesting to assess whether reduced inhibitory GSK3 phosphorylation is a common feature in ALS patients, and whether the UNC13A genetic status affects this.

Altogether, we tested a hypothesis generated by a high-throughput genetic screen, performing multiple steps of validation. We did this in a variety of model systems with different mutations, because factors that rescue among multiple systems are more likely to be applicable to sALS patients, the vast majority of the disease population. With this work, we showed that PP2A and GSK3 are modifiers of FUS toxicity in *Drosophila* and patient sMNs. Our results are the first to demonstrate that PP2A modifies FUS-mediated toxicity *in vivo*, and provide a better understanding as to how GSK3 activity may be regulated in ALS. Furthermore, we unraveled a novel mechanistic link between PP2A, GSK3 and kinesin-1, which may lead to mitochondrial transport defects in FUS-ALS (Fig. 9). Therefore, this study has a strong impact on understanding the underlying mechanisms of FUS-ALS pathogenesis.

## Methods

### *Drosophila* lines and maintenance

*Drosophila melanogaster* strains were maintained on standard medium (62.5g/L cornmeal, 25 g/L yeast, 7g/L agar, 16.9 g/L dextrose, 37.5ml/L golden syrup, 9.375 ml/L propionic acid, 1.4g/L hydroxybenzoate, 14.0 ml/L ethanol) in a 12h light/dark rhythm. FUS lines have been described previously^9^. The following stocks were obtained from the Bloomington *Drosophila* Stock Center (BDSC): D42-Gal4 (8816), Gal80-ts (7019), nSyb-Gal4 (51635), sgg RNAi#1 (31308), sgg RNAi#2 (31309), sgg RNAi#3 (38293), sgg RNAi#4 (35364), *UAS-sgg WT* (5361), *UAS-sgg S9A* (5255), null mutant mts^XE2258^ (5684), null mutant sgg (4095). The following lines were obtained from the Vienna *Drosophila* RNAi center (VDRC): mts RNAi#1 (35171), mts RNAi#2 (35172), mts RNAi#3 (41924). The Khc line *w;;p[Khc+]* was provided by William Saxton (University of California, Santa Cruz).

For backcrossing: The *w1118* (Canton-S10) line was used as a background control, for experiments using TRiP lines a stock consisting of W+v-X chromosome in the *w1118* genetic background (provided by Teresa Niccoli, University College London) was used. Males of the indicated genotypes were crossed to virgins of the background stock for 1 generation, before virgins were selected and crossed into the background stock for 6 further generations. Flies were rebalanced using balancers crossed into the background stock for 6 generations.

### Genetic screen of *Drosophila*

The BDSC deficiency kit was ordered from the Bloomington Drosophila Stock Center. Deficiencies covering chromosomes 2, 3, 4 were crossed to flies of the following genotype: w*;;UAS FUS(R521G), D42-Gal4 / TM6BGal80* and reared at 25℃. Pharate TM6B-negative pupae were moved to a petri dish and followed for 72h. The percentage of eclosed flies was defined as ratio of the number of empty pupal cases to the number of total pupal cases.

### Eclosion phenotype with modifying lines

Prior to experiments, sgg (31308, 31309, 38293, 35364) and mts (35171/GD, 35172/GD, 41924/GD) RNAi lines were backcrossed to *W+v-w1118* (TRiP lines, marked with vermillion) or *w1118* (VDRC lines, marked with miniwhite) control for six generations, to ensure that results are not affected by different genetic backgrounds of the flies. 5 sgg or mts RNAi virgins were crossed to 3 *w;;D42-Gal4, UAS-FUS/TM6BGal80* males and females were allowed to lay eggs for 48h. Parents were removed from the vials and the progeny were allowed to grow at 25C. 14 days after set up, eclosion was scored, counting the number of eclosed VS total TM6B-negative pupae. *w1118* and *W+v-* background strains were crossed in as controls.

### Eclosion phenotype with LiCl and OA

Lithium chloride (LiCl) (0-20mM, L9650, Sigma Aldrich) or okadaic acid (OA) (0-50nM, 459620, Sigma Aldrich) was dissolved in milliQ H_2_O and added to standard food at the indicated concentrations. For the 0 condition, H_2_O alone was added. w-;;UAS-FUS WT and R521G males were crossed to D42-Gal4 virgins. Females were allowed to lay on drug containing food for 48h. Crosses were left to develop at 25 °C and eclosion was scored 14 days after set up.

### Lifespan

The indicated lines were crossed to *w;Gal80ts;D42-Gal4, UAS-FUS/TM6B* males on grape agar plates supplemented with yeast paste. Approximately 50 virgins and 20 males were used per cross. Eggs were collected after 24h into PBS and seeded at a standard density into bottles. Bottles were allowed to develop for 21 days at 18℃. Adult male flies (female in the case of sgg null mutant crosses) were briefly anaesthetized on CO_2_ and split into vials at a density of 10 flies per vial. Approximately 150 flies (15 vials) were analyzed per condition, exact N numbers are given in supplementary material. Flies were tipped onto fresh food twice a week, and deaths were scored every other day at the beginning and every day towards the end of the experiment. Statistical significance was assessed using log-rank test.

### iPSC lines

A previously characterized *FUS* mutant iPSC line from a 17-year old male ALS patient carrying a *de novo* mutation (P525L) was used^3,4^. The FUS-ALS line was compared with its isogenic CRISPR-Cas9 gene-edited isogenic control (P525P) generated by CellSystems (Troisdorf, Germany)^3^. Cells were cultured and differentiated into MNs according to a well-established protocol as described^3^. The cells were previously collected from the donor with the approval of the ethical committee of the University Hospitals Leuven (S50354).

### Immunofluorescence of iPSC-derived MNs and FUS mislocalization

Drugs (LiCl L9650 Sigma Aldrich, tideglusib SML0339 Sigma Aldrich, OA 459620 Sigma Aldrich) were prepared fresh and dissolved in water (DMSO for tideglusib). Cells were mock treated using an equivalent volume of vehicle. On day30 of differentiation, cells were briefly washed with PBS, and fixed with 4% paraformaldehyde (PFA) in PBS for 15min at room temperature. Cells were then washed 3×10min with PBS, followed by a blocking step for 1h with 5% normal donkey serum (NDS) (D9663, Sigma Aldrich) in 0.1% PBS-Triton X-100 (PBS-T). Cells were incubated with primary antibodies overnight at 4℃ in 2% NDS-PBS-T (Table S5). The following day, cells were washed 3×5min with 0.1% PBS-T and the secondary antibodies (Table S6) were added in 2% NDS-PBS-T for 1h at room temperature. Cells were then washed 2x with PBS and NucBlue^TM^ Live ReadyProbes^TM^ Reagent (Invitrogen) was added at a concentration of 2 drops/mL PBS for 20min at room temperature to stain the nuclei (Hoechst 33342). Cells were washed 2x with PBS and coverslips were mounted onto slides with ProLong Gold Antifade mounting reagent (P36934, Thermo Fisher Scientific). Cells were imaged with an inverted confocal microscope (SP8 DMi8, Leica Microsystems). Captured images were analyzed using ImageJ software. For nucleocytoplasmic quantification, regions of interest (ROIs) corresponding to the nucleus of the cell and the soma were drawn using the signal in the Hoechst channel before measurements in the FUS channel were taken. A nearby non-cellular ROI was used to determine the average background of each ROI, and these values were subtracted from the measurements. The ratio of the background subtracted raw integrated densities was used to calculate the nucleocytoplasmic ratio.

### NMJ analysis

The same FUS mutant and isogenic control iPSC-derived MNs as described above were used. In addition, human myoblasts were isolated from a biopsy from a 55-year old healthy woman and cultured as described previously^31,78^. On day10 motor neuron neuro-progenitor cells (MN-NPCs) were seeded in the two wells and the channel on one side of the microgrooves in the microfluidic device (Xona Microfluidics, XC150) at 125,000 cells per well (250,000 NPCs/device). Similarly, myoblasts were seeded in the two wells and the channel opposite to the MNs in the device at 20,000 cells per well (40,000 myoblasts/device). Myoblasts were differentiated into myotubes and MN-NPCs into sMNs following the established protocol^31^. On day18, a chemotactic and volumetric gradient was established. MN compartments received 100 μl/well neuronal medium without neurotrophic factor, and myotube compartments received 200 μl/well neuronal medium supplemented with 10 ng/mL BDNF (PeproTech, Rocky Hill, NJ, USA, cat. no. 450-02), GDNF (PeproTech, cat. no. 450-10) and CNTF (PeproTech, cat. no. 450-13) in addition to 20 mg/mL laminin (Sigma-Aldrich, cat. no. L2020-1MG) and 0.01 mg/mL recombinant human agrin protein (R&D Systems, cat. no. 6624-AG-050). On the same day, both sides of the device were treated with a drug (LiCl 1mM 48h, tideglusib 15μM 48h, okadaic acid 1nM 72h). Mock co-culture devices were kept in parallel without drug treatments. The growth factor and volume gradients, including agrin/laminin, were maintained at each medium change, which was performed every other day for 10 days. On day28, devices were fixed and stained, and images were taken using an inverted SP8 DMi8 Leica confocal microscope. Quantification of NMJ formation was manual and blinded, based on the neurite / presynaptic marker morphology and/or based on colocalization between presynaptic marker neurofilament heavy chain/synaptophysin with post-synaptic marker α-bungarotoxin in myosin-heavy chain-labelled multinucleated myotubes.

### SH-SY5Y cell cultures

SH-SY5Y cells (94030304, Sigma) were cultured in T175 flasks in DMEM:F12 Glutamax medium (ThermoFisher Scientific) supplemented with 10% Fetal Bovine Serum (FBS). Media were changed twice a week. For this, cells were washed with Versene (15040066, ThermoFisher Scientific) and lifted with 0.05% trypsin (25300054, ThermoFisher Scientific). Cells were maintained in an incubator (37℃, 5% CO_2_). For western blotting, cells were seeded in 6-well plates at a density of 0.15×10^6^ cells/mL. 48h after splitting, fresh media were added containing drug or vehicle. Treatment lasted for 24h.

### SH-SY5Y cell lysis for western blotting

24h after treatment, cells were washed briefly with PBS. Cells were lysed on ice in RIPA buffer (R0278-500ML, Sigma-Aldrich) supplemented with protease inhibitor (cOmplete^TM^, EDTA-free protease inhibitor cocktail (1836170001, Sigma-Aldrich) and phosphatase inhibitor (Phos-STOP^TM^ 4906837001, Sigma Aldrich). Cells were scraped and the lysate was pipetted into a pre-chilled Eppendorf tube. Lysis was allowed to proceed for 20min on ice and cells were centrifuged at 16000xG for 10min at 4℃. Supernatant was transferred into a fresh tube and Pierce BCA Protein assay was performed (23225) according to the manufacturer’s instructions. Samples were brought to equal concentrations using addition of RIPA buffer, and Pierce^TM^ Lane Marker Reducing Sample Buffer (39000, Thermo Fisher Scientific) was added to a final concentration of 1X. The samples were boiled for 5min at 95℃.

### Motor neuron lysis for western blotting

Cells were briefly washed with DPBS. Cells were scraped in DPBS and transferred to a pre-chilled Eppendorf tube. They were centrifuged for 5min at 1000xG at 4℃. Pellets were then dissolved in RIPA buffer with 1× Phos-STOP (Roche) and 1× cOmplete EDTA-free protease inhibitor (Roche). Lysis was allowed on ice for 30min. Samples were centrifuged at full speed for 10min at 4℃ and the supernatant was collected. Pierce BCA Protein assay (23225) was performed. Samples were brought to equal concentrations, supplemented with Pierce^TM^ Lane Marker Reducing Sample Buffer (39000, Thermo Fisher Scientific), to a final concentration of 1X and boiled for 5min at 95℃.

### *Drosophila* head lysate for western blotting

At the indicated age, adult flies were frozen in liquid nitrogen. Heads were removed by vigorously banging the tube containing frozen flies. 10 fly heads were collected per condition on dry ice and before being homogenized at room temperature in 100ul of 2x Pierce^TM^ Lane Marker Reducing Sample Buffer (39000) using a power pestle. Samples were boiled in 95℃ for 5min and centrifuged at 20000xG at room temperature for 5min and the supernatant moved to a fresh tube.

### Purification of KLC1p Antibody

Generation of the rabbit antibody to KLC1 phosphorylated on serine-460 was described by Mórotz et al. 2019. Prior to use, the antibody was purified using the phospho-peptide (CKVDSphosPTVTTTLKNL), which was synthesized by GenScript, using the High-Affinity Antibody Purification Kit (L00404, GenScript) following the manufacturer’s protocol.

### Phosphorylation assays in HEK293T cells

HEK293T cells were transfected with FLAG-tagged WT and P525L FUS and treated with 10nM calicheamicin γ1 (MedChemExpress, HY-19609) for 2 h. Next the cells were treated with 10nM OA (dissolved in H_2_O) for different time periods. Vehicle-treated cells were treated with the same amount of H_2_O. Cells were scraped on ice and western blot analysis was performed.

### Western blotting

Western blot was performed using NUPAGE 4-12% Bis-Tris 1.0mm Mini Protein Gels (ThermoFisher Scientific). The PAGE-ruler prestained protein ladder was used as reference (26616, ThermoFisher Scientific). After electrophoresis (140V, 400mA), the gel was transferred to a PVDF membrane (IPVH00010 Immobilon-P transfer membrane, Sigma-Aldrich), using the Mini Trans-Blot Cell system (BIORAD). The membrane was blocked in 5% non-fat dry milk (9999S, Cell Signaling) or 5% BSA (11930, SERVA) diluted in Tris-buffered saline with Tween-20 (TBS-T) for 1h at room temperature. The membrane was then incubated with primary antibody at 4℃ overnight in 5% BSA in TBS-T. The following day, after washing 3×10min with TBS-T, secondary antibody was added in 5% BSA in TBS-T for 1h at room temperature. Details of all primary and secondary antibodies are given in Tables S5 and S6. The membrane was then washed 3×10min in TBS-T and developed with the Pierce^TM^ ECL Western Blotting Substrate (Pierce ECL, 32106, Thermo Fisher Scientific) or the SuperSignal™ West Pico PLUS Chemiluminescent Substrate (34580). Images were taken using a chemiluminescence instrument (ImageQuant LAS4000).

### Live cell imaging of mitochondrial transport and tracking analysis

To measure mitochondrial neurite transport, iPSC-derived sMNs at differentiation day30 were incubated with 50nM MitoTracker^TM^ Green FM (M7514, Invitrogen) in neuronal medium for 20min at 37℃. After 20min, cells were washed and incubated in BrainPhys^TM^ Imaging Optimized Medium (05796, STEMCELL Technologies) for imaging. Images were taken using the Operetta CLS High-Content Analysis System (PerkinElmer) with a 40x objective at 37℃ and 5% CO_2_. The MitoTracker^TM^ Green was excited at ∼495nm and 1-second time-lapse images were taken for 200 seconds. Video files were analyzed with ImageJ using TrackMate v3.8.0 plugin for total mitochondria quantification and tracking, and time/distance kymographs to quantify the number of moving mitochondria in a selected neurite segment. Moving mitochondria are represented by tilted lines, whereas stationary mitochondria can be discerned as straight vertical lines.

### Statistics

Data are presented as mean ± SEM, unless indicated otherwise. Statistical analyses were performed in GraphPad Prism 9.

## Data availability

All data generated or analyzed during this study are available from the authors.

## Acknowledgments

We are grateful to Lindsey Robyns and Martine Noblet for preparation of the *Drosophila* media. We thank Valerie Bercier for comments on the paper. We would like to thank Gabor M. Morotz and Christopher CJ Miller for providing the KLC1 S460p antibody serum and advice on the purification of the antibody. We thank the following investigators for providing us with *Drosophila* stocks: Teresa Niccoli, University College London; William Saxton, University of California, Santa Cruz. The kinesin heavy chain, head region (Khc) antibody developed by J.M. Scholey, University of California, Davis was obtained from the Developmental Studies Hybridoma Bank, created by the NICHD of the NIH and maintained at The University of Iowa, Department of Biology, Iowa City, IA 52242. Stocks obtained from the BDSC (NIH P40OD018537) were used in this study. Transgenic fly stocks were obtained from the VDRC. We thank somersault18:24 for valuable assistance with Fig.9 assembly. This work was supported by VIB, KU Leuven (C1 and “Opening the Future” Fund), the “Fund for Scientific Research Flanders” (FWO-Vlaanderen), the Thierry Latran Foundation, the “Association Belge contre les Maladies neuro-Musculaires – aide à la recherché ASBL” (ABMM), the Muscular Dystrophy Association (MDA), the ALS Liga België (A Cure for ALS), Target ALS and the ALS Association (ALSA). P.T. is funded by a PhD Fellowship of FWO-Vlaanderen (11C7623N). T.M. is supported by an FWO postdoctoral fellowship (1246821N) and an EU Horizon 2020 Programme for Research and Innovation Marie Sklodowska-Curie actions (grant 845692). P.V.D. holds a clinical investigatorship of the FWO and is supported by the E. von Behring Chair for Neuromuscular and Neurodegenerative Disorders, the Fund ‘Een Hart voor ALS’ and the ‘Laevers Fund for ALS Research’. L.V.D.B. is supported by the Generet Award for Rare Diseases.

## Author contributions

P.T. and T.M. designed the experiments. P.T. performed most of the experiments, did the data analysis and wrote the manuscript. J.S. performed the blots of FigS3 and the *Drosophila* screen. W.S. assisted with experiments and fly maintenance. A.S.C. assisted with the iPSC-sMNs cultures and provided instructions on how to perform the mitochondrial transport analysis. K.S.D. and A.M.B.C. assisted with the NMJ experiments and analysis. A.H. and A.P. helped to perform mitochondrial transport experiments. P.V.D. provided the human material and guidance designing experiments. T.M. and L.V.D.B. supervised the project, discussed the results and edited the manuscript. All authors read and approved the final version of the manuscript.

---

**Fig. S1.**
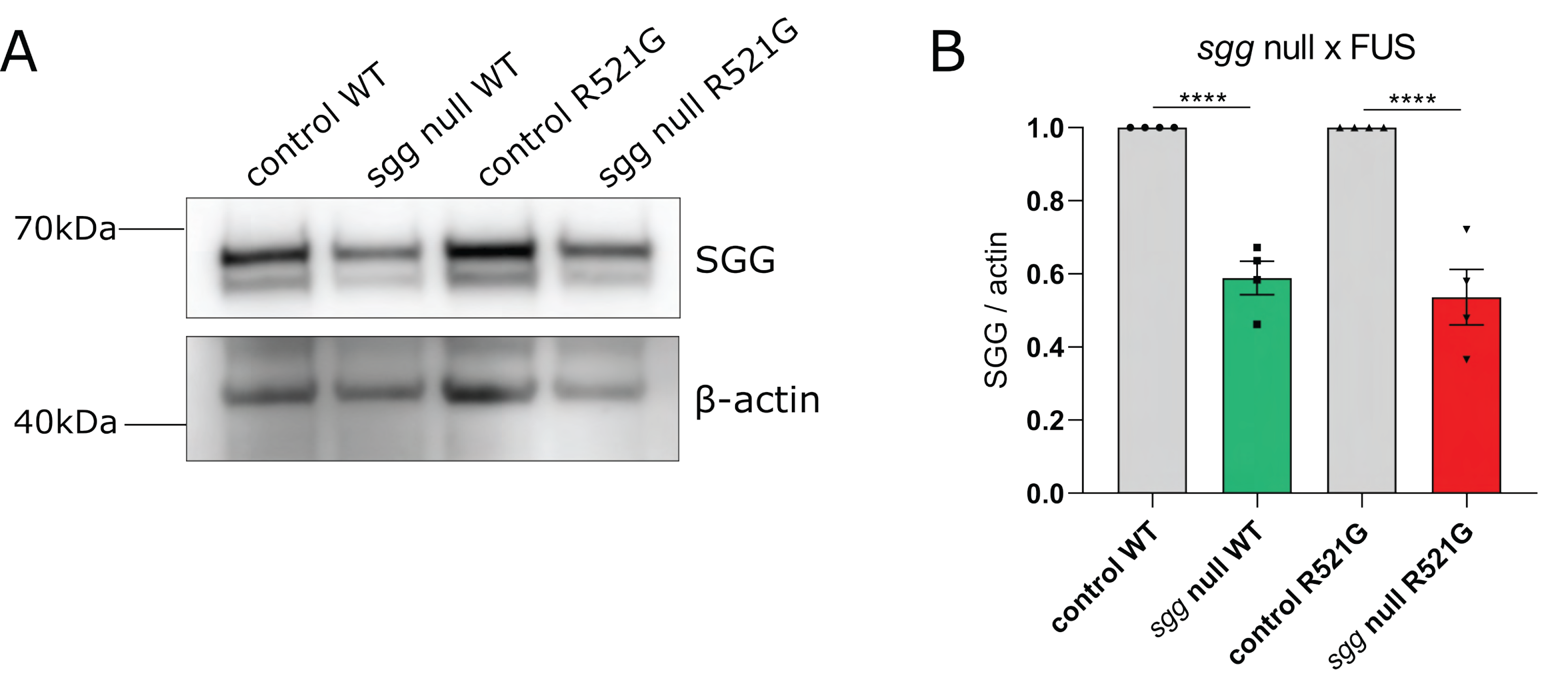
sgg null mutant reduces the levels of sgg protein by ∼50%. **A**. Western blot for sgg protein levels in sgg null mutant line. Beta-actin serves as loading control. (N=3) **B**. Quantification of panel A shows that the sgg null mutant line induces ∼50% reduction of sgg protein levels. (N=3, mean ± SEM, one-way ANOVA with Sidak’s multiple comparisons test) ****p< 0.0001

**Fig. S2.**
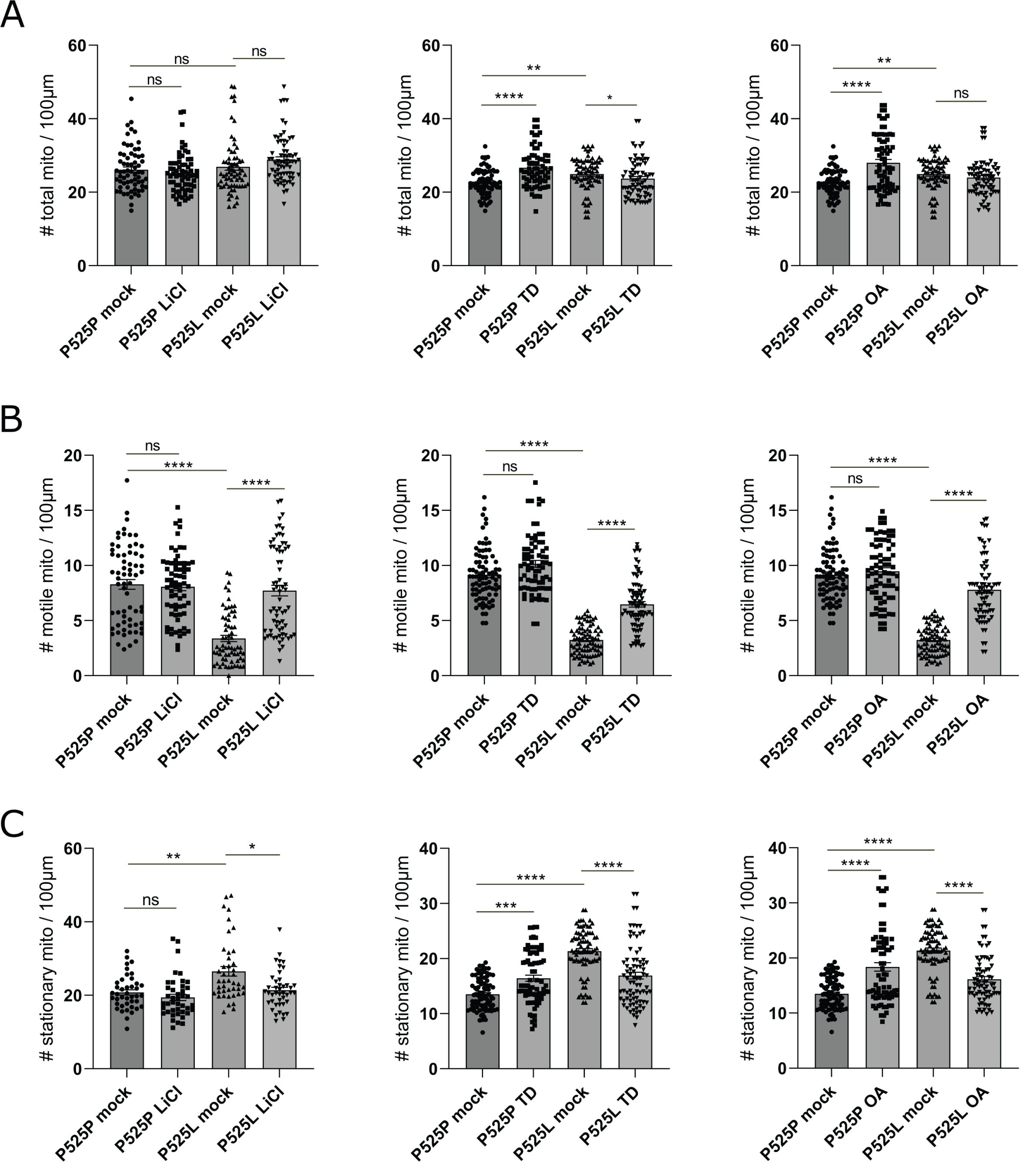
Additional data on the effect of LiCl, TD and OA on mitochondrial transport in sMNs. **A to C.** Quantification of total (A), motile (B) and stationary (C) mitochondria normalized to 100μm neurite length. Total mitochondria are calculated as the sum of motile and stationary events. Data are represented as mean ± SEM, N=3 independent differentiations, Kruskal-Wallis with Dunn’s multiple comparisons test. ****p< 0.0001, ***p<0.001, **p<0.01, *p<0.05, ns= not significant

**Fig. S3.**
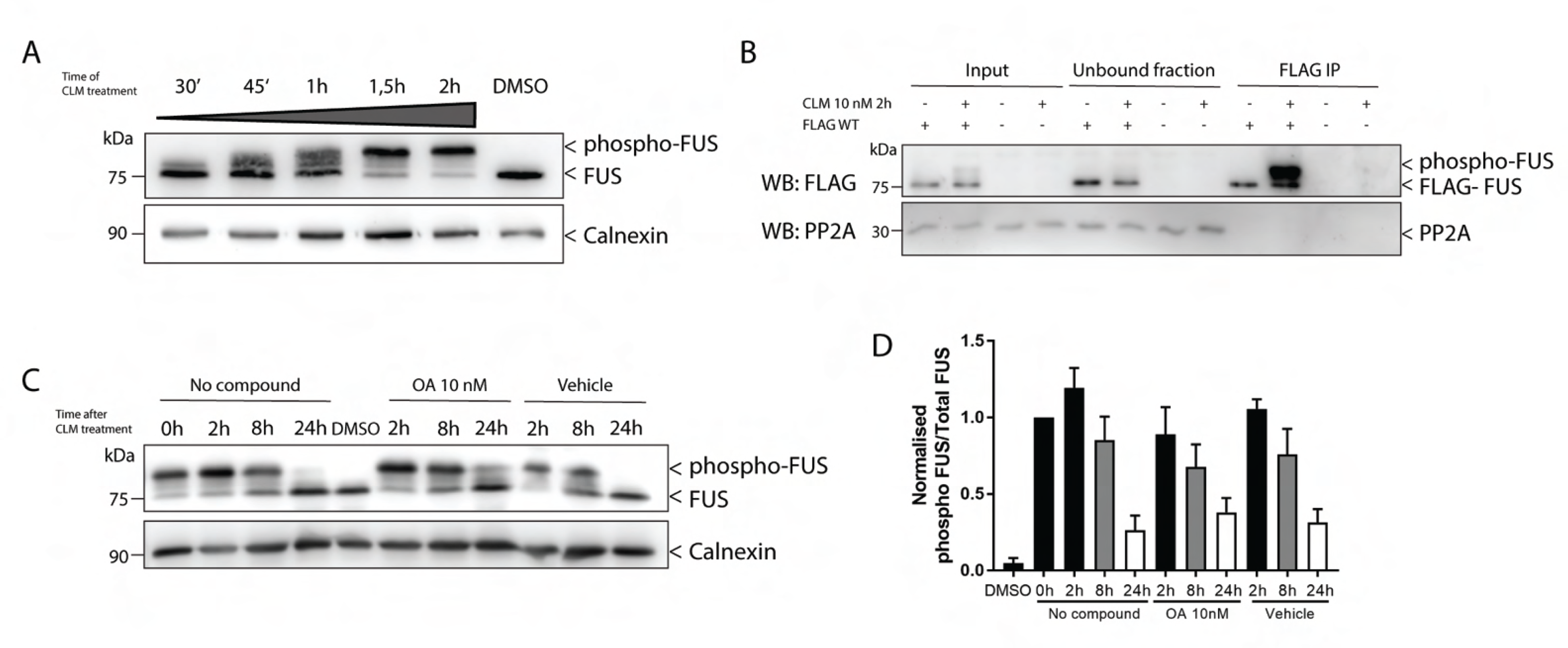
PP2A is an upstream suppressor of FUS toxicity. **A.** FUS phosphorylation by 10nM CLM treatment is examined at different time points. After 45 min of CLM treatment, phospho-FUS was observed and after 2 h of CLM treatment, endogenous FUS was almost completely phosphorylated. **B.** Lysates from FLAG-WT hFUS-expressing HEK293T cells treated with 10nM CLM were immunoprecipitated using an anti-FLAG antibody. No interaction between FUS and PP2A was observed. **C.** HEK293T cells treated with 10nM CLM showed almost complete dephosphorylation after 24 h (no compound). PP2A inhibition with 10nM OA could not affect the dephosphorylation rates of FUS. **D.** Quantification of Western blot of panel C.

**Table S1:** Overview of the candidate modifiers that resulted from the genetic screen.

|  | Modifying gene | CG number | Human orthologue | Rescue |  | RNAi lines tested | RNAi lines positive | Null Mutants |  | Null mutants tested | Null mutants positive |
| --- | --- | --- | --- | --- | --- | --- | --- | --- | --- | --- | --- |
|  |  |  |  | Female D42/R521G (%) | Male D42/R521G (%) |  |  | Female D42/R521G (%) | Male D42/R521G(%) |  |  |
| Unbiased approach | NaCh | CG8178 | SCNN1G | 9 | - | 1 | 1 |  |  |  |  |
|  | CG34458 | CG34458 | CSTG | - | 4 | 1 | 1 |  |  |  |  |
|  | Herp | CG14536 | HERPUD2 | 1 | 3 | 2 | 2 | - | - | 2 | 0 |
|  | Mts | CG7109 | PPP2CA | 8 | 1 | 1 | 1 | 12 | 0 | 4 | 2 |
|  | CG7231 | CG7231 | FAM151B | 0 | 3 | 2 | 1 | 7 | 0 | 2 | 1 |
|  | Sirup | CG7224 | SDHAF4 | 5 | 2 | 1 | 1 | 1 | 0 | 1 | 0 |
|  | R2d2 | CG7138 | STAU2 | 4 | 4 | 2 | 1 | 4 | 0 | 2 | 1 |
|  | SmE | CG18591 | SNRPE | 5 | 5 | 2 | 1 | 4 | 0 | 1 | 1 |
|  | CG31915 | CG31915 | COLGALT1 | 1 | 0 | 2 | 1 | 85 | 40 | 1 | 1 |
|  | CG7277 | CG7277 | COQ6 | 0 | 7 | 1 | 1 | 22 | 0 | 1 | 1 |
|  | qkr54B | CG4816 | KHDRBS1 | 1 | - | 2 | 1 | 55 | 27 | 1 | 1 |
|  | NT5E-2 | CG30104 | NTE | 1,4 | - | 2 | 1 |  |  | 2 | 0 |
|  | Patronin | CG33130 | CAMSAP2 | 13 | 13 | 2 | 1 | 17 | 4 | 3 | 1 |
|  | CG6568 | CG6568 | unknown | 1 | - | 1 | 1 | 3 | 2 | 1 | 1 |
|  | l(2)k01209 | CG4798 | UCKL1 | 3 | - | 1 | 1 | - | - | 3 | 0 |
|  | CG18467 | CG18467 | GID8 | 2 | - | 1 | 1 |  |  |  |  |
|  | FMR1 | CG6203 | FMR1 | 3 | 3 | 5 | 3 | 5 | 6 | 2 | 2 |
|  | Sgg | CG2621 | GSK3B | 63 | 45 | 5 | 2 | 7 | - | 3 | 1 |
| Literature | HRB87F | CG12749 | HNRNPA2B1 | 2 | - | 1 | 1 | 4 | - | 1 | 1 |
|  | HRB27C | CG10377 | DAZAP1 | 3 | 6 | 2 | 2 | 50 | 64 | 2 | 1 |
|  | Parp | CG40411 | PARP1 | 17 | 4 | 2 | 1 |  |  |  |  |
|  | Sgg | CG2621 | GSK3B | 63 | 45 | 5 | 2 | 7 | - | 3 | 1 |
|  | qkr54B | CG4816 | KHDRBS1 | 1 | - | 2 | 1 | 55 | 27 | 1 | 1 |
|  | FMR1 | CG6203 | FMR1 | 3 | 3 | 5 | 3 | 5 | 6 | 2 | 2 |

**Table S2:** The statistical information of the RNAi lifespans (from Fig.2).

|  | Experiment | Condition | N<br>(number of flies) | Median lifespan<br>(days) | P value |
| --- | --- | --- | --- | --- | --- |
| sggRNAi x FUS | WT FUS | control | 152 | 29.5 | - |
|  |  | RNAi #1 | 149 | 36.0 | 5.24807E-58 |
|  |  | RNAi #2 | 151 | 34.0 | 3.62347E-30 |
|  |  | RNAi #3 | 155 | 36.0 | 5.57358E-39 |
|  |  | RNAi #4 | 156 | 36.0 | 2.92332E-54 |
|  | R521G FUS | control | 160 | 28.0 | - |
|  |  | RNAi #1 | 148 | 36.0 | 3.37108E-64 |
|  |  | RNAi #2 | 149 | 36.0 | 4.90037E-60 |
|  |  | RNAi #3 | 148 | 36.0 | 1.99196E-60 |
|  |  | RNAi #4 | 146 | 36.0 | 6.91336E-65 |
| mtsRNAi x FUS | WT FUS | control | 142 | 31.5 | - |
|  |  | RNAi #1 | 134 | 38.5 | 3.00091E-48 |
|  |  | RNAi #2 | 126 | 34.0 | 3.87413E-17 |
|  |  | RNAi #3 | 139 | 34.0 | 1.09269E-18 |
|  | R521G FUS | control | 158 | 29.5 | - |
|  |  | RNAi #1 | 128 | 38.5 | 1.05996E-52 |
|  |  | RNAi #2 | 139 | 38.5 | 1.50108E-56 |
|  |  | RNAi #3 | 128 | 31.5 | 3.0097E-38 |

**Table S3:**
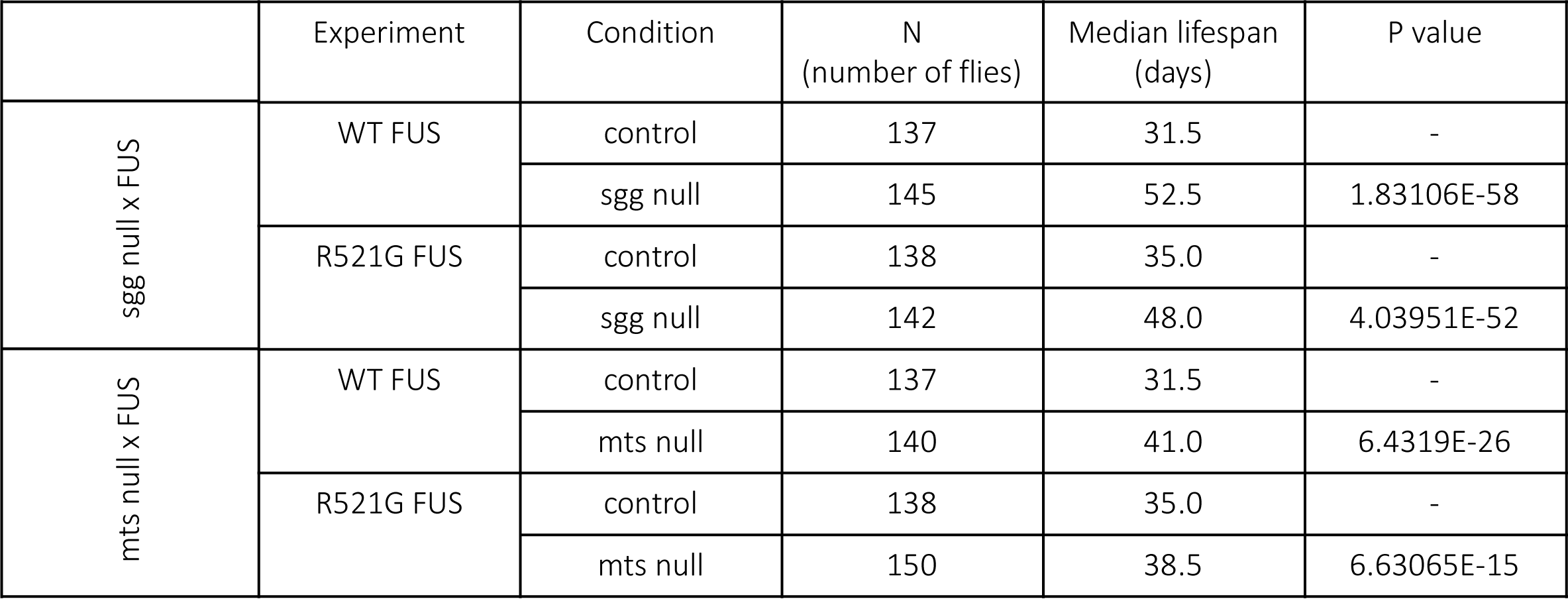
The statistical information of the null mutant lifespans (from Fig.2).

|  | Experiment | Condition | N<br>(number of flies) | Median lifespan<br>(days) | P value |
| --- | --- | --- | --- | --- | --- |
| sgg null x FUS | WT FUS | control | 137 | 31.5 | - |
|  |  | sgg null | 145 | 52.5 | 1.83106E-58 |
|  | R521G FUS | control | 138 | 35.0 | - |
|  |  | sgg null | 142 | 48.0 | 4.03951E-52 |
| mts null x FUS | WT FUS | control | 137 | 31.5 | - |
|  |  | mts null | 140 | 41.0 | 6.4319E-26 |
|  | R521G FUS | control | 138 | 35.0 | - |
|  |  | mts null | 150 | 38.5 | 6.63065E-15 |

**Table S4:**
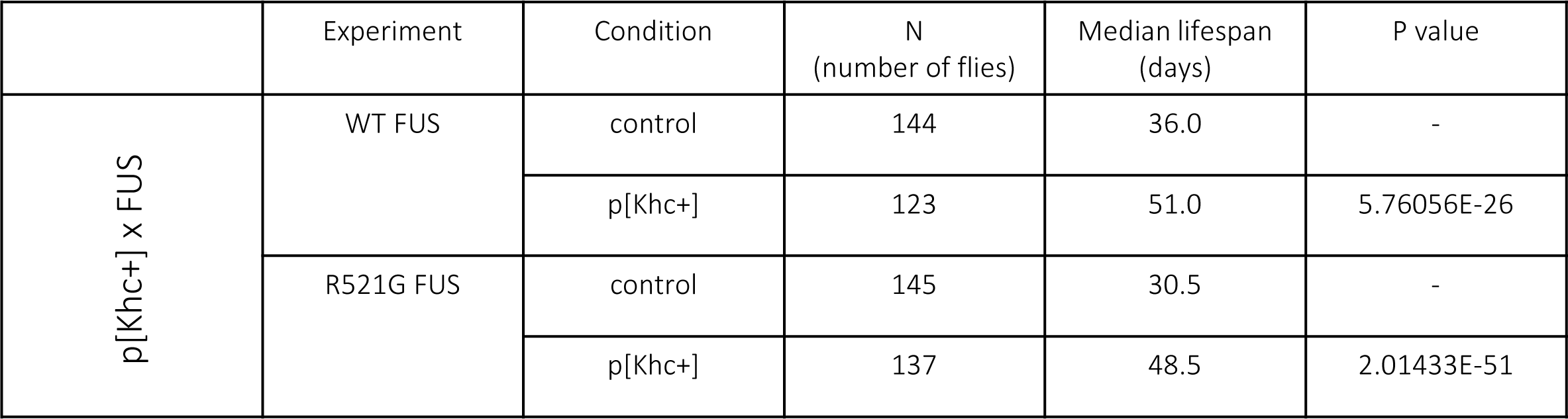
The statistical information of p[Khc+] lifespans (from Fig.8).

**Table S5:**
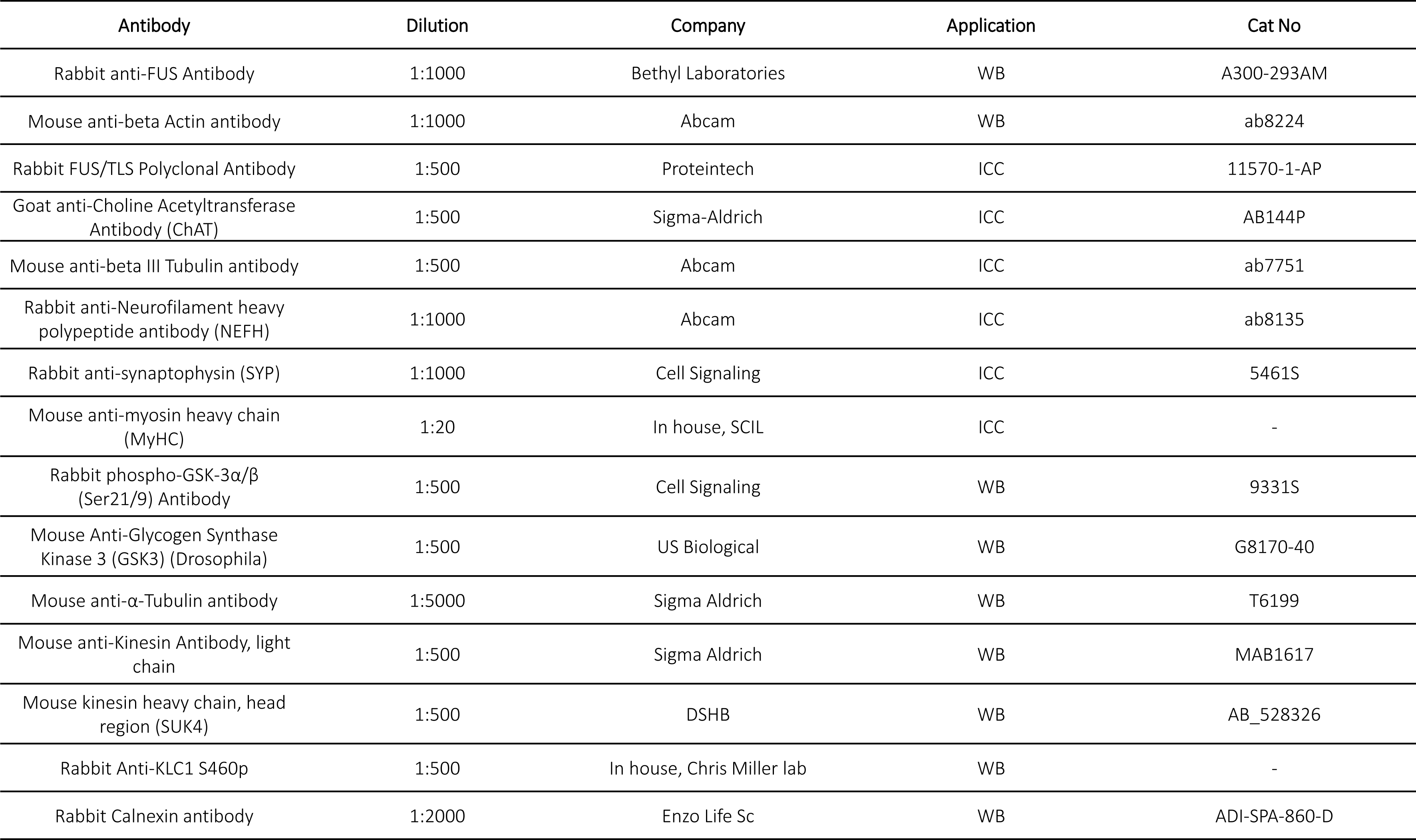
Primary antibody overview.

**Table S6:**
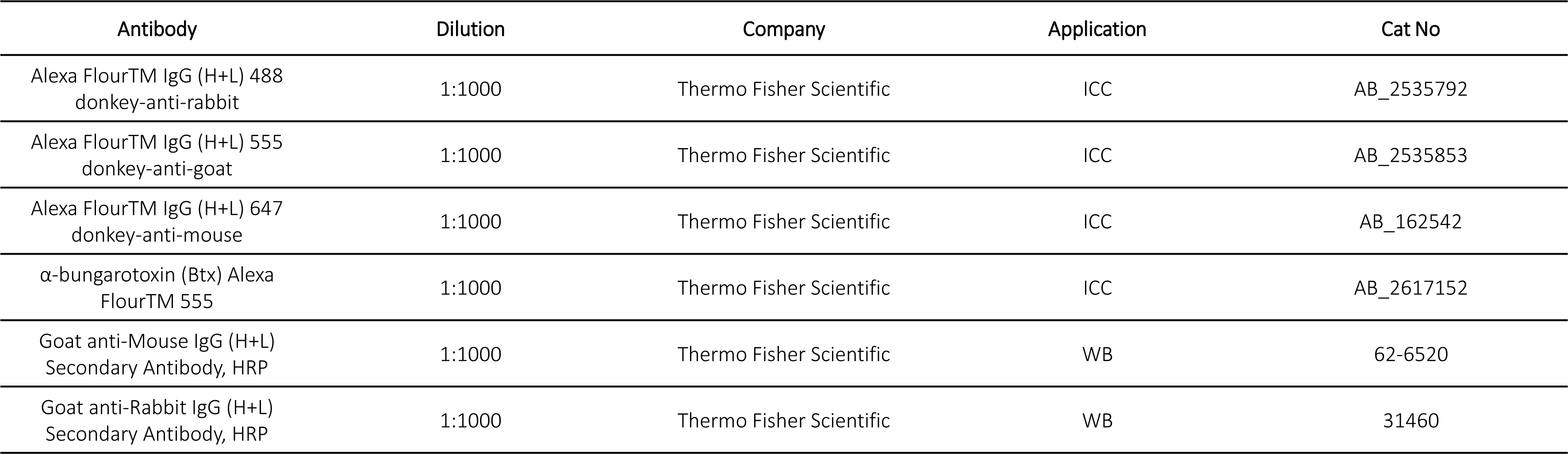
Secondary antibody overview.

